# Identifying win-win opportunities and trade-offs for sustainable agriculture to improve agricultural productivity and soil carbon sequestration: A global meta-analysis

**DOI:** 10.1101/2025.09.02.673637

**Authors:** David G. Encarnation, Robert S. Powell, Pete Smith, Adam F. A. Pellegrini

## Abstract

Faced with accelerating climate change and growing food insecurity, sustainable agriculture has the potential to address both issues by enhancing soil carbon sequestration and crop yields. However, it remains unclear whether and when sustainable agricultural practices create win-wins or trade-offs between soil carbon sequestration and food production, and how these outcomes vary across crop types, practice combinations, and environmental conditions. We conducted a global meta-analysis of 2,975 paired yield and topsoil carbon observations (and 498 subsoil carbon observations) from 487 articles to assess the simultaneous impacts of reduced tillage, cover cropping, complex crop rotations, and crop residue retention on soil organic carbon stocks and maize, wheat, rice, and soybean yields. Our analysis revealed that sustainable agricultural practices generally enhanced soil carbon and crop yields, although with considerable heterogeneity. Individual practices often resulted in management trade-offs, as different practices (and crop-practice combinations) optimized either yields or soil carbon sequestration. However, practice stacking often alleviated these trade-offs by combining complementary practices. Win-wins were most likely in marginal agricultural lands characterized by warmer temperatures, water limitation, and low nitrogen application rates – conditions that represent a substantial proportion of global agricultural land area. However, our analysis revealed that the subsoil carbon responses to sustainable agricultural practices were highly variable and may offset topsoil gains, which may limit effective policy design and large-scale implementation.

## 1.0 Introduction

Modified agricultural practices have the potential to both mitigate emissions from the food system (which accounts for a third of anthropogenic emissions) (Crippa et al., 2021) and improve the food supply for over 2 billion people facing food insecurity (FAO, 2024). Given that land available for agriculture is limited, a key opportunity is addressing both issues by adopting practices that simultaneously mitigate climate change and increase food production. One pathway is improving soil quality, namely *via* increases in soil organic carbon (SOC) (Smith, 2012). SOC is often correlated with greater yields of several staple crops (Ma et al., 2023; Oldfield et al., 2019), and increasing soil carbon sequestration in agricultural lands may contribute 47% of climate mitigation potential for agriculture; but range from 0.25-6.78 Gt CO_2_-eq yr^−1^ (Roe et al., 2019). Thus, there is scope to achieve both goals of increasing food production and sequestering carbon simultaneously.

Agricultural practices thought to improve SOC and yield include reduced tillage, cover cropping, complex crop rotations, and improved crop residue management (Rehberger et al., 2023). Adopting these practices can affect crop yields and soil carbon sequestration, in both positive (Bai et al., 2019; Lessmann et al., 2022; Lobell et al., 2024; McClelland et al., 2021; Ogle et al., 2019; Qiu et al., 2024; Xia et al., 2018; Zhao et al., 2022) and negative (Lobell et al., 2024; Ogle et al., 2019; Pittelkow et al., 2015) ways. This creates opportunities to identify win-win scenarios for food production and climate change mitigation but also requires an understanding of where there are trade-offs. Indeed, recent modelling work has suggested that these trade-offs may be more common than previously thought (Abdalla et al., 2019; Lobell et al., 2024; McClelland et al., 2025; Qiu et al., 2024; Sun et al., 2020). But empirical studies investigating the simultaneous impacts of these practices on soil carbon and crop yields have been rare and region-specific (e.g., winter cover cropping in the Argentinean Pampas (Alvarez et al., 2017). Accordingly, relating soil carbon and yield responses to changes in agricultural management remains one of the main barriers in estimating the climate mitigation potential and scaling implementation of sustainable practices (Amelung et al., 2020; Oldfield et al., 2024).

Comparing effects across multiple practices and crop types is crucial to maximize gains. Combining multiple practices may help to alleviate trade-offs (Abdalla et al., 2019; McClelland et al., 2025; Pittelkow et al., 2015; Sun et al., 2020) or increase the gains in yields and/or soil carbon stocks (Qiu et al., 2024), though they may also reduce benefits relative to the individual practices (e.g., Prairie et al., 2023).

The magnitude and distribution of win-wins and trade-offs also depend on agronomic conditions and environmental constraints. While there is substantial heterogeneity in the relationships between yields and SOC responses and agronomic and environmental gradients, some trends have been identified. Most of the practices perform best in warmer conditions, where enhanced decomposition converts carbon inputs from residue retention or cover crops to soil organic matter, or where reduced tillage counteracts enhanced decomposition (Poeplau & Don, 2015; Prairie et al., 2023; Shang et al., 2021). Practices like reduced tillage and residue retention that may help conserve soil moisture often perform best in drier, water-limited regions (Pittelkow et al., 2015; Sun et al., 2020), though other practices like cover cropping or complex crop rotations that increase water consumption may perform better in moist areas (Abdalla et al., 2019). Similarly, soil properties can influence crop productivity and place constraints on SOC accrual. Practices like residue retention and reduced tillage may be more likely to improve yields in sandier soils (Xia et al., 2018) that are water-limited, and SOC benefits tend to be greater in soils with low initial SOC concentrations (Liu et al., 2014; Rehberger et al., 2023). Recent work suggests that the saturation of mineral-associated organic carbon (MAOC) impacts the accrual and vulnerability of SOC stocks and offers a more refined estimate of SOC sequestration potential (Georgiou et al., 2025). Lastly, agronomic decisions like fertilizer use can produce different responses: cover cropping and complex crop rotations tend to increase yields more in low-N systems (Abdalla et al., 2019; King & Blesh, 2018), whereas other practices may perform better in high-N conditions (Shang et al., 2021; Xia et al., 2018). It is also expected that the SOC benefits from these practices take time to accumulate (Liu et al., 2014; Poeplau & Don, 2015; Prairie et al., 2023; Xia et al., 2018) but may saturate (Shang et al., 2021). While the effects of these environmental and agronomic gradients on crop yields and soil carbon outcomes are well-documented (often with competing trends), there has not been an assessment of multiple sustainable practices in multiple crops on combined yield and SOC changes that would facilitate practice comparisons aimed at developing optimal practices.

Moreover, previous research on the effects of these practices on soil carbon sequestration have often been limited to the topsoil (King et al., 2024; McClelland et al., 2021; Poeplau & Don, 2015; Sun et al., 2020). Studies that have investigated subsoils have shown both neutral (Prairie et al., 2023; Qiu et al., 2024) and negative (e.g., Ogle et al., 2019) effects, which has thus far limited the scientific consensus around whether subsoil carbon responses may offset topsoil SOC increases (Oldfield et al., 2024; Paustian et al., 2016). Similarly, limited global-scale research suggests that practices such as residue retention and reduced tillage may increase microbial biomass carbon (Li et al., 2018), which may have broader soil health and emissions implications.

We use a global meta-analysis to investigate the potential of four agricultural practices – improved crop residue management, reduced tillage, cover cropping, and complex crop rotations – to sequester carbon in the soil while maintaining or increasing yields of four staple crops: maize, wheat, rice, and soybeans.

## 2.0 Methods

### 2.1 Data Collection

#### 2.1.1 Literature Search

We comprehensively searched Web of Science and SCOPUS for studies that quantified both the yield and soil carbon impacts of four agricultural practices: cover cropping, complex crop rotations, reduced tillage, and crop residue retention (Table S1). We searched for studies that investigated the impact of any of these practices on soil carbon concentrations or stocks **and** yields of four main staple crops (maize, wheat, rice, and soybeans) (Lists S1 & S2). The search included all English-language primary articles published until November 17, 2022. After duplicates were removed, this yielded 13,945 unique articles. We present a detailed description in the Supplementary Information of study selection criteria.

#### 2.1.2 Data Compilation & Management

In total, we identified 487 papers published between 1978 and 2024 that met our inclusion criteria (Figure S1). A complete list of included publications is in the Supplementary Information. The practices investigated are defined in Table S1. Cover cropping and complex crop rotations are often hard to distinguish because the distinction is often based on whether the “cover” crop is harvested or not, which is not always reported. As such, we often combined the two practices into cropping system intensification (“Intensification”), following Prairie et al (2023). We did not differentiate between different seasons of crops (e.g., spring wheat vs winter wheat; early vs late-season rice) as papers did not reliably report this information.

For each study, we collected the means, number of replications (sample size) and estimates of variability (standard deviation, standard error, coefficient of variation) for the SOC and yield parameters in control (conventional) and treatment (sustainable) groups. We imputed the missing standard deviations using the coefficient of variation for all observations with reported standard deviations and then used this average coefficient of variation to impute the missing standard deviations (imputation methods are robust (Furukawa et al., 2006; Pellegrini et al., 2023)).

As we wanted to compare soil carbon stocks, which represent the actual amount of carbon stored (rather than soil carbon concentrations), we occasionally imputed missing bulk density values. Instead of using global gridded datasets, which would give the same values for control and treatment groups, we used the mean bulk density values for conventional separately from sustainable observations (see Supplementary Information for greater detail). This preserves the effect of agricultural practices on bulk density which has been shown to be significant (e.g., Al-Kaisi & Lal, 2020; Liu et al., 2014). We also aggregated soil carbon stocks across depth using bulk densities to first calculate stocks in each layer (Prairie et al., 2023). We used a threshold of 30 cm to separate between topsoil (0-30 cm) and subsoil (>30 cm) layers (see Supplementary Information for details on details on overlapping layer depths). This resulted in 2975 aggregated topsoil observations, 498 aggregated subsoil observations, and 531 aggregated full profile observations, with maximum depths in the full profile ranging from 35-105 cm. We also collected 326 aggregated topsoil observations of microbial biomass carbon concentrations.

Environmental and agronomic covariate data were either taken from the papers or supplemented using global gridded datasets (Table S2). Data were taken from spatially explicit global datasets using latitude and longitude provided by the study or based on nearest landmark (e.g., city).

### 2.2 Meta-analysis

#### 2.2.1 Effect size calculation

All statistics were run in R v4.3.2 (R Core Team, 2023). The natural log of the response ratio (RR) was used as the effect size to measure the effect of the sustainable agricultural practices on soil carbon stocks and crop yields. The response ratio numerator is the ‘sustainable’ practice and the denominator is the ‘conventional’ practice. Thus, a value > 0 illustrates a gain in SOC or yield. Effect sizes were calculated with the *escalc* function from the *metafor* package (Viechtbauer, 2010). We weighted observations by the inverse of the variance. SOC and yield response ratio outliers were identified based on having a z-score > 3 and were removed before analysis.

#### 2.2.2 Variable Importance

To identify important predictors of soil carbon and yield responses, we first removed highly correlated continuous variables (Pearson correlation coefficient ≥ 0.70). We then used the R package *metaforest* (Lissa, 2020) to fit meta-regression models, which are robust to overfitting and non-linear relationships and incorporates meta-analytical variance and weights.

We performed recursive variable pre-selection for both the complete dataset and the practice-specific data subsets (see Supplementary Information for details on subset definitions) in *metaforest* with 10,000 iterations and 100 replications using the *preselect* function, retaining variables that improved model performance in >90% of iterations. We then optimized the tuning hyperparameters (Table S3) using the *train* function within the *caret* package (Kuhn, 2008) and selected top models to minimize root-mean-square error (RMSE). The practice-specific metaforest models were fitted with different numbers of moderators depending on practice-specific variables (e.g., cover crop functional group). Important predictors were identified using an importance threshold of 0.7 in each top model.

#### 2.2.3 Univariate models and meta-regressions

To visualize how the soil carbon and yield responses were distributed across different thresholds of environmental and agronomic variables, we fitted univariate mixed effects models using the *rma.mv* function in the *metafor* package. This approach allowed us to account for potential non-independencies of data by including study sites as a random effect.

After establishing variable importance using *metaforest*, we visualized the individual effects of important predictors on topsoil SOC and yield responses using meta-regression analysis. For each continuous predictor variable, we used the *predict* function from the *metafor* package to generate prediction curves by varying the focal predictor across its observed range while holding all other important variables constant at their mean values (or mode for categorical variables). This allowed for interpretation of the effects of individual predictors while accounting for other important predictors. We transformed response ratios into percentage change for visualization.

#### 2.2.4 Model assumptions

We assessed the presence of publication bias using funnel plots, Egger’s regression tests (Egger et al., 1997), the trim-and-fill method (Duval & Tweedie, 2000), and Fail-Safe N Analysis (Rosenberg, 2005). See Supplementary Information for more detail.

## 3.0 Results

The complete dataset spanned 403 study sites spanning all six continents with arable agriculture, with 2,975 observations of paired yield and topsoil organic carbon stock measurements (Figure S2). The representativeness of the dataset in terms of global distributions of important environmental covariates is shown in Figures S3 and S4.

### 3.1 Practices improve both yields and topsoil soil carbon stocks globally

Overall, the practices were beneficial for topsoil carbon and crop yields. Win-wins— significant increases in both yield and topsoil SOC—were present in 18.1% of observations, and roughly twice the number of observations (37.5%) contained significant gains in either one of SOC stocks or crop yields under improved management. In contrast, it was rare (5.7% of observations) to experience significant losses of SOC or yield, and exceptionally rare (1% of observations) to experience losses in both soil carbon and yields (Figure 1). Observations without significant effects totalled 37.8%. Thus, while there was substantial variability in the magnitude of effects, the majority (55.6% of observations) demonstrated significant gains in yield and/or topsoil SOC, while significant losses occurred in only 6.7% of observations.

**Figure 1.**
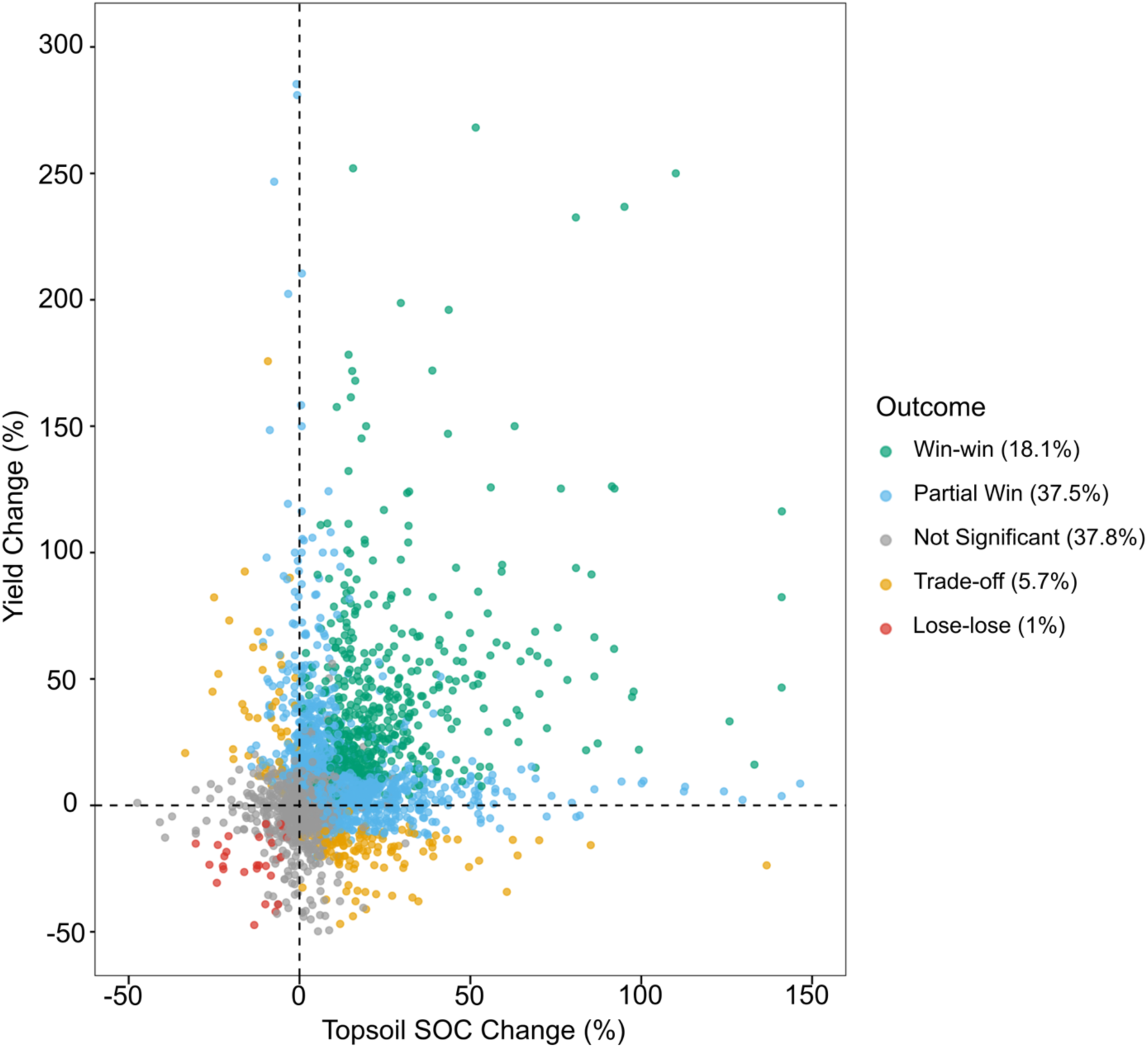
Distribution of yield and topsoil soil carbon stock responses. Points illustrate the percentage change in crop yields and topsoil SOC for each aggregated topsoil observation. Colours indicate the significance based on the 95% confidence interval of each point estimate (estimated using 1.96 * standard error): Win-win = both responses significantly positive. Partial Win = One response significantly positive while the other response is neutral. Not Significant = both responses non-significant. Trade-off = one response is significantly negative. Lose-lose = both responses significantly negative.

### 3.2 Magnitude of topsoil carbon and yield responses depend on crop type and type of practice implemented

Across all four staple crops investigated, sustainable agricultural practices consistently increased mean topsoil SOC stocks by approximately +10.0% (Figure 2a). The yield response was more variable. Significant yield increases were observed in maize and wheat systems (average increase of +13.1% and +8.2%, respectively). Soybean yields increased by an average of +12.9%, but this was not significant, nor was the +5.7% average increase in rice yields. Thus, maize and wheat exhibit more consistent gains in yields while all exhibit consistent gains in topsoil SOC.

**Figure 2.**
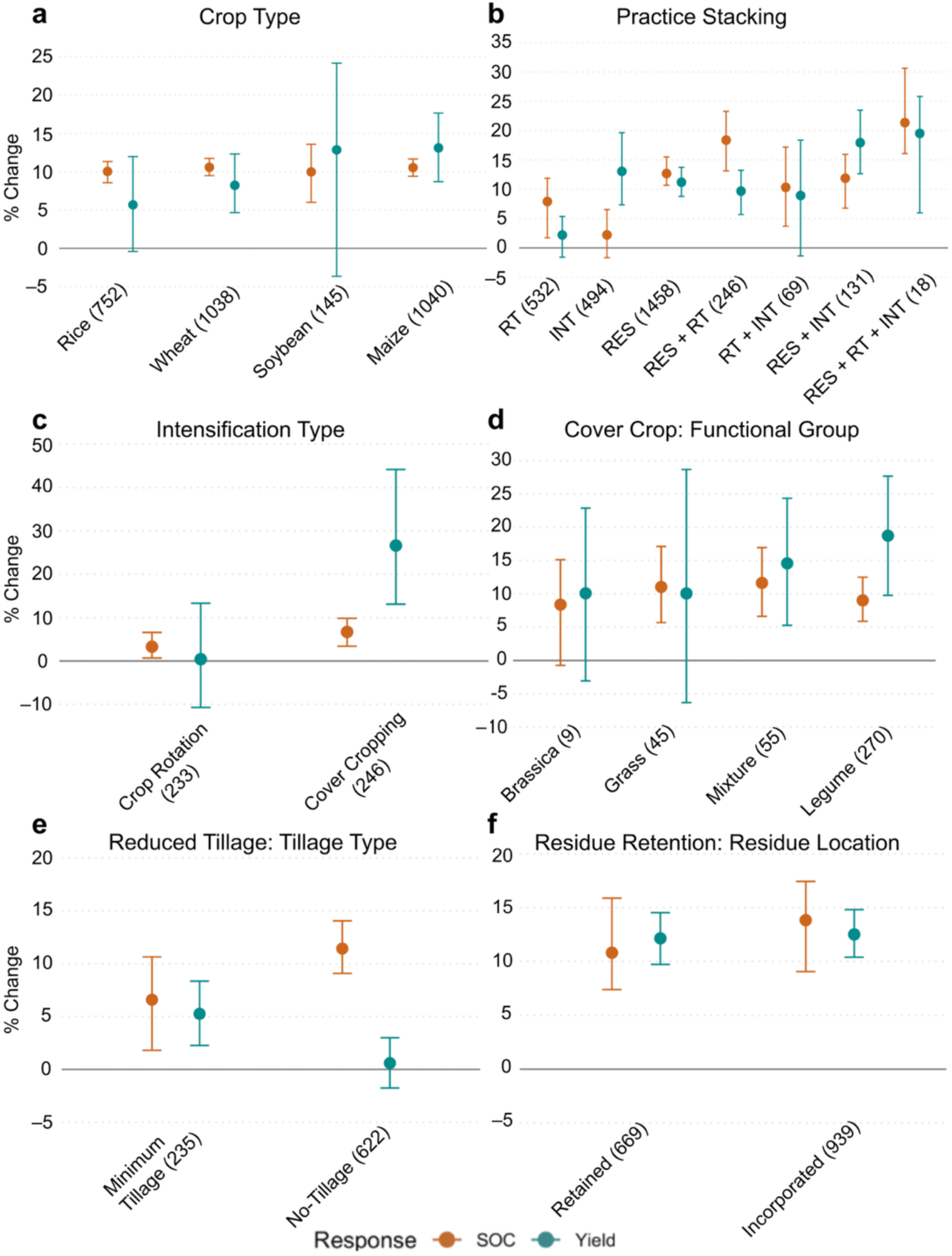
Overall topsoil SOC and crop yield responses. Broken down by crop type (**a**) and combination of practices used (**b**). **a** and **b** use the full dataset and c-f use practice-specific datasets for cover cropping (c), crop rotations (d), reduced tillage (e), and residue retention (f). All analyses based on univariate mixed effects models. Points represent mean estimates and error bars represent bootstrapped 95% confidence intervals, such that if the bars do not overlap the x-axis then the effects are significant. Values in parentheses represent sample sizes. In b, RT = reduced tillage, INT = cropping system intensification (cover cropping and/or crop rotations) and RES = residue retention. In c-f, practice-specific data subsets were used, which include any observation where a practice is adopted, even when there is practice stacking (e.g., the Reduced Tillage subset will include observations where the treatment is both reduced tillage and residue retention). See Supplementary Information for more details of practice-specific subsets and bootstrapping procedures.

Of the individual practices investigated in isolation, reduced tillage by itself led to sizeable topsoil SOC gains (+7.9%) yet insignificant yield increases. On the other hand, intensification (cover cropping and/or crop rotations) by itself led to large average yield increases (+13.1%) but insignificant SOC increases (Figure 2b). Within intensification, cover cropping alone slightly increased SOC (+6.7%) and led to a large significant increase in crop yields (+26.6%), while crop rotations alone slightly increased SOC (+3.3%) and had no effect on crop yields (+0.4%, p > 0.5) (Figure 2c). Residue retention by itself produced large gains in both SOC (+12.7%) and yields (+11.2%) (Figure 2b). Thus, while all individual practices are beneficial for at least one outcome, management trade-offs may arise given the variability of SOC and yield responses to different practices.

### 3.3 Practice implementation methods influenced the magnitude of topsoil soil carbon and yield responses

Each individual practice has multiple methods of implementation, which were important for predicting topsoil SOC and crop yield responses. For example, the implementation of reduced tillage can be broadly divided into minimum tillage and no-tillage methods (Table S1), which affected the magnitude and significance of SOC and yield changes: Switching from conventional to no-tillage led to a larger SOC increase (+11.4%) than minimum tillage (+6.6%); but no-tillage did not affect crop yields (Figure 2e) whereas minimum tillage led to increases (+5.3%).

Cropping system intensification can be partitioned into cover cropping and crop rotation methods (Figure 2c), as well as the species used for cover cropping. The SOC response was consistently positive in all cover crop functional groups except *Brassica spp.* and was marginally higher when using grasses (+11.0%) compared to legumes (+9.0%), though this difference was not significant. Conversely, using a legume cover crop led to a significantly positive yield response (+18.7%) whereas using a grass cover crop species did not significantly affect crop yields. Using a mixture of cover crop functional groups led to large and significant increases in both SOC (+11.6%) and crop yields (+14.6%) (Figure 2d).

Residue retention effects depended on the depth of addition: the SOC response was greater when crop residues were incorporated (+13.8%) into the soil rather than retained on the soil surface (+10.8%). Despite this, both residue incorporation and surface retention led to similar yield responses (+12.5% and +12.1%, respectively) (Figure 2f).

Thus, even within a practice, the method of implementation can change not only the magnitude and significance of yield and topsoil SOC changes, but whether the practice generates co-benefits between SOC and yield.

### 3.4 Stacking practices enhanced gains in topsoil SOC and yields

Combining multiple practices can have synergistic effects that often exceeded the benefits of individual practices. Combining other practices with reduced tillage tended to increase the average topsoil SOC response (from +12.7% to +18.4% for residue retention and from +2.2% to +10.3% for intensification) though it did slightly decrease the average yield response (from +11.2% to +9.7% for residue retention and from +13.1% to +8.9% for intensification). Combining other practices with intensification tended to increase the average yield response (from +11.2% to +17.9% for residue retention and from +2.2% to +8.9% for reduced tillage, though the latter two were not statistically significant). Residue retention was the most broadly beneficial practice and combining other practices with residue retention increased both SOC and yield responses. This may have helped overcome the negligible impacts of intensification on SOC (increased the average response from +2.2% to +11.9%) and reduced tillage on yields (increased the average response from +2.2% to +9.7%). Lastly, though the sample size was small (*n* = 18), combining all three practices led to the largest mean increases in SOC (+20.4%) and crop yields (+19.5%) (Figure 2b). Thus, stacking multiple practices can improve outcomes and alleviate neutral or negative impacts of individual practices.

### 3.5 Efficacy of practices varied by crop type

Crop-practice interactions were important determinants of the topsoil SOC and yield responses. In rice systems, all individual practices increased topsoil SOC, though the average response was largest with residue retention (+11.3%), followed by reduced tillage (+9.9%) and intensification (+6.2%). Intensification and residue retention significantly increased rice yields (+15.4% and +8.7%, respectively), and reduced tillage did not significantly affect yields. In soybean systems, residue retention increased topsoil SOC (+9.9%) but reduced tillage and intensification did not have significant effects. Conversely, intensification increased soybean yields (+13.0%) but reduced tillage and residue retention did not. In wheat systems, all practices increased topsoil SOC, though the average response was largest with residue retention (+12.8%), followed by reduced tillage (+8.8%) and intensification (+5.3%). Residue retention also increased maize yields (+9.7%), but reduced tillage and intensification did not have significant effects. Lastly, in maize systems, residue retention significantly increased topsoil SOC (+13.8%) and yields (+15.2%). Intensification significantly increased maize yields (+13.1%) but had no effect on SOC, and reduced tillage did not significantly impact either outcome. These findings demonstrate that the effect of the practices may vary by crop type, but there is not always a clear ‘best practice’ for a given crop.

### 3.6 SOC and yield gains were greater on marginal land with historically less intensive management

Several climatic, soil, and farm management gradients were significantly related to topsoil SOC and yield responses. For climate, the greatest SOC response was observed in arid regions (+17.7%), though it was consistently positive across humid, dry sub-humid, and semi-arid zones (+9.6%, +12.6%, and +9.4%, respectively). Conversely, the lowest yield response was observed in arid regions (+4.2%) relative to wetter zones (+8.6%, +11.3% and +10.6%, in humid, dry sub-humid, and semi-arid regions, respectively) (Figure 3a). Thus, while SOC and yields may increase across all aridity zones, more arid regions are less likely to experience yield gains despite the large increase in SOC stocks.

**Figure 3.**
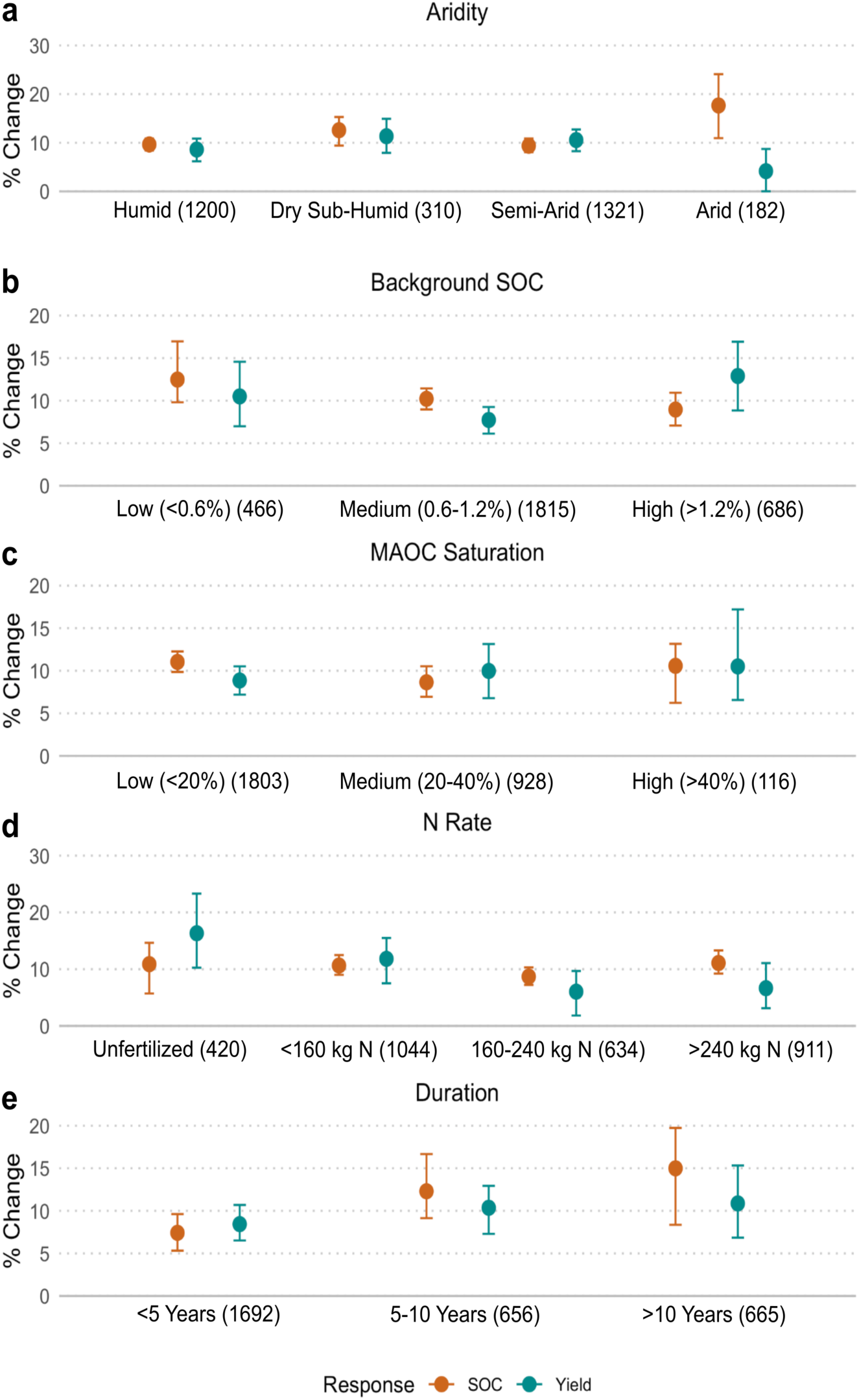
Overall topsoil SOC and crop yield responses broken down by selected soil, climate and management predictors. All analyses based on univariate mixed effects models. Points represent mean estimates and error bars represent bootstrapped 95% confidence intervals, such that if bars do not overlap the x-axis then the effects are significant. Values in parentheses represent sample sizes. Aridity index (AI) derived from Zomer et al. (2022). Arid and hyper arid: 0 < AI < 0.2; Semi-Arid: 0.2 ≤ AI < 0.5; Dry sub-humid: 0.5 ≤ AI <0.65; Humid: AI ≥ 0.65. Background SOC derived from HWSD (FAO & IIASA, 2023). MAOC Saturation is derived from Georgiou et al. (2022) and refers to the realized amount of mineral-associated organic carbon (MOC) divided by the potential amount of mineral-associated carbon (MOC_max_). Nitrogen application rate and study duration derived from the studies themselves.

For soil properties, the greatest SOC response was observed in soils with low initial (background) SOC contents (+12.5%), followed by medium- and high-SOC content soils (+10.2% and +9%, respectively). But the background SOC content had a variable impact on the yield response, with the greatest yield response in high-SOC soils (+12.9%) and the lowest yield response in medium-SOC soils (+7.7%) (Figure 3b). Similarly, the average SOC response was greatest (albeit marginally) in the soils with the lowest level of mineral-associated organic carbon (MAOC) saturation (+11%) whereas the average yield response was greatest in the soils with the highest level of MAOC saturation (+10.5%) (Figure 3c). Thus, soil carbon gains tend to be larger in soils with low initial soil carbon concentrations, while the trend with MAOC saturation is more variable.

Nitrogen application rate did not have a clear effect on the SOC response, with average responses ranging from +8.7% in the 160-240 kg N/ha category to 11.1% in the >240 kg N/ha category. There was, however, a clear negative relationship between nitrogen application rate and yield response. The greatest yield response was observed in unfertilized systems (+16.3%), followed by low-N systems (<160 kg N/ha; +10.6%), and finally medium-N (+6%) and high-N systems (+6.6%) (Figure 3d). Thus, nitrogen application reduces the relative gains in yield but the gains in SOC are consistent across different application levels.

Lastly, both SOC and yield responses accumulated over time. The greatest SOC response was in long-term studies (+15%; >10 years), followed by medium-term (+12.3%; 5-10 years) and smallest in short-term studies (+7.4%; <5 years). Similarly, the yield response was greatest in long-term and medium-term studies (+10.9% and +10.4%, respectively) and smallest in short-term studies (+8.4%) (Figure 3e). Thus, both SOC and yield responses may be maximized over longer time periods, but the effect of time may have a stronger impact on the SOC response.

### 3.7 SOC and yield responses across practices were greater in stressful and low N environments

The effects of reduced tillage on both topsoil SOC and yield responses were primarily influenced by climatic variables (Table S8) and were positively associated with mean annual temperature (Figure 4a), potential evapotranspiration (Figure 4b) and aridity (negatively associated with the aridity index which is the inverse i.e., more arid regions have a lower aridity index) (Figure 4c). The mean yield response became negative in areas with lower mean annual temperature (<10° C) and in humid climates (aridity index > 0.65), whereas the SOC response was consistently positive and changed minimally across the gradients. Thus, reduced tillage maximizes the combination of SOC and yield responses in warmer and drier conditions.

**Figure 4.**
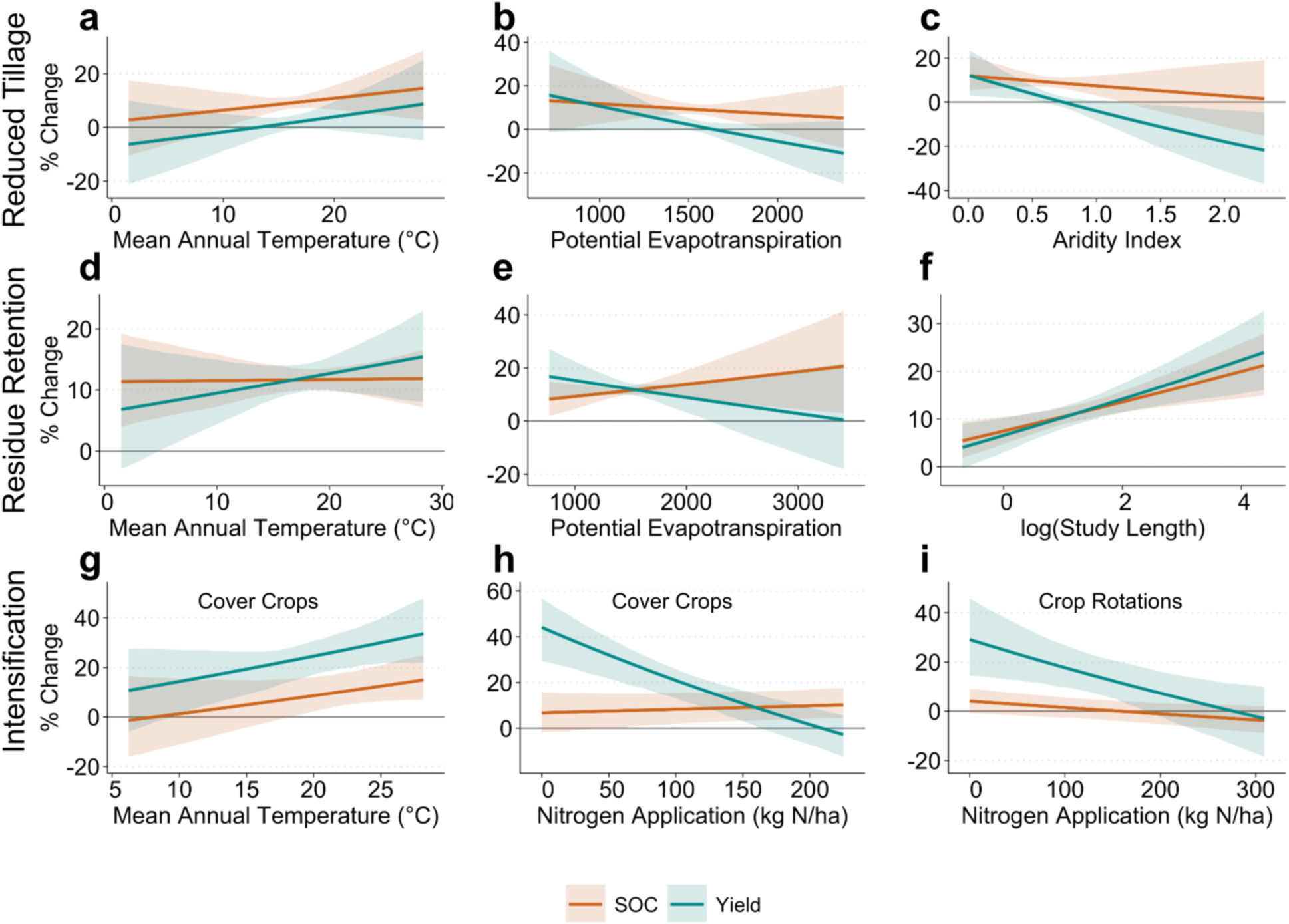
Practice-specific partial regressions. Models derived from important predictors from practice-specific SOC and yield sub-models. Partial regressions generated by varying one predictor while holding others at their means and categorical variables at their most common level when applicable. Cover cropping plots (4g & 4h) are set to the most common crop type (Rice) and crop rotation plot (4i) is set to the most common management level (Intensification alone). Prediction curves based on 100 equally spaced model predictions (see Supplementary Information). Error bars represent bootstrapped 95% confidence intervals, such that if bars do not overlap the x-axis then the effects are significant. Note that aridity index is the inverse of aridity i.e., more arid regions have a lower aridity index.

The effects of intensification practices (cover cropping and crop rotations) were influenced primarily by farm management variables such as crop type and N application rate, and to a lesser extent by climatic conditions (Table S8). For cover cropping, topsoil SOC and yield responses were positively associated with mean annual temperature (Figure 4g). Unlike reduced tillage, the yield response to cover cropping was consistently positive, and the SOC response only became significantly positive in warmer climates (MAT >20° C).

The yield responses to both cover cropping and crop rotations declined as the nitrogen application rate increased (Figures 4h,i), though the thresholds at which the yield response became positive varied by crop type (Figure S5). Rice and maize yields increased when nitrogen application was below 160 kg N/ha, wheat yields increased with nitrogen application less than 100 kg N/ha, and soybean yields never significantly increased across the entire nitrogen gradient.

The effects of residue retention were influenced by both climatic and management variables (Table S8). The yield response increased with increasing mean annual temperature (Figure 4d) though the SOC response was unaffected. The yield response correlated negatively but the SOC response correlated positively with potential evapotranspiration (Figure 4e). Both SOC and yield responses were positively associated with study duration (Figure 4f). Thus, the benefits of residue retention take time to accrue despite some climatically controlled trade-offs when trying to maximize for either SOC or yields.

Taken together across all practices, these results suggest that sustainable agricultural practices tend to perform best (and thus win-wins are more likely) in stressful environments such as warmer, more arid and less-fertile systems. Increased nitrogen application tends to reduce the efficacy of intensification practices, especially for yields.

### 3.8 Topsoil SOC changes were associated with increased microbial biomass

Microbial biomass carbon concentrations increased across all four practice combinations with sufficient data. Reduced tillage had the largest effect of any individual practice (Figure 5a) especially under no-tillage (+34.8%) rather than minimum tillage (+23.8%) (though the confidence intervals overlapped) (Figure 5b). Intensification (cover cropping and crop rotations) also increased MBC (+17.3%), followed by residue retention (+12.6%). The effect of residue retention on MBC was greater when residue was retained on the soil surface (+30.6%) rather than incorporated into the soil (+14.0%), in contrast with topsoil SOC gains, but the confidence intervals of the two methods overlapped (Figure 2f; Figure 5c). Aridity was the most important predictor of the microbial biomass carbon response (Table S10), with the greatest average MBC response in dry sub-humid regions (+25.8%) and the smallest mean response in semi-arid regions (+19.3%). Thus, while microbial biomass carbon concentrations tend to increase across the board, different practices (and within-practice implementation methods) affect the magnitude of the changes.

**Figure 5.**
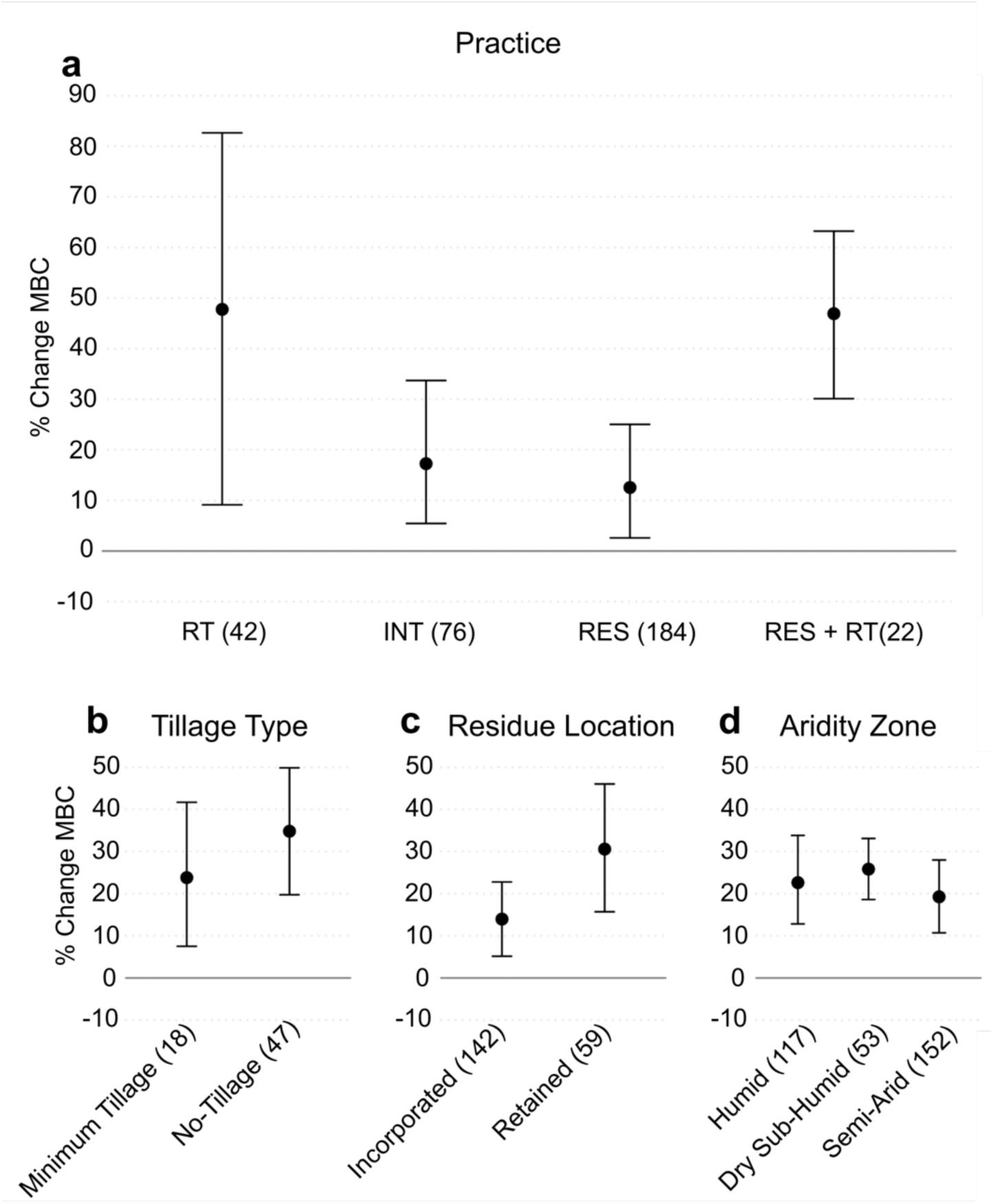
Microbial biomass carbon (MBC) responses to sustainable practices. Broken down by type of practice used (**a**), type of tillage in the reduced tillage subset (**b**), residue location in residue retention subset (**c**), and aridity zone (**d**). All analyses based on univariate mixed effects models. Points represent mean estimates and error bars represent bootstrapped 95% confidence intervals, such that if the bars do not overlap the x-axis then the effects are significant. Values in parentheses represent sample sizes. In a, RT = reduced tillage, INT = cropping system intensification (cover cropping and/or crop rotations) and RES = residue retention.

### 3.9 Subsoil SOC responses were more variable and may have offset topsoil SOC gains

While all practices apart from intensification had significantly positive effects on topsoil SOC (Figure 2b), the subsoil effects were much more variable. We were only able to assess these responses on a subset of our data (*n_topsoil_* = 2,975 vs *n_subsoil_* = 498), and thus part of the variation observed in the subsoil effects may be due to (i) which studies were included and (ii) lower confidence because of smaller sample sizes.

Only residue retention had a significantly positive effect on subsoil SOC stocks (+43.7%), though the estimates varied widely (95% CI [+4.0%, +82.8%]). The remaining practices did not significantly affect subsoil SOC stocks, though the average effect of intensification was slightly negative (-7.3%) and the average effect of reduced tillage on subsoil SOC stocks was substantially negative (-24.3%).

When we aggregated topsoil and subsoil values to investigate the response of the full soil profile, reduced tillage no longer had a significantly positive effect on SOC stocks (p > 0.50), indicating that the increases in the topsoil were offset by decreases in the subsoil. The only individual practice with a significantly positive effect on total SOC stocks was residue retention, where moderate topsoil SOC increases (+12.7%) were bolstered by large average increases in the subsoil (+43.7%) to produce a +20.5% increase across the whole soil profile. Although sample sizes were small, combining residue retention with reduced tillage (RES + RT) significantly increased total SOC stocks (+17.2%), as did combining residue retention with intensification (RES + INT; +12.9%). Taken together, while subsoil responses were highly variable, considering subsoil SOC changes produced different responses depending on the practice. Reduced tillage and intensification practices produced negative mean effects on subsoil SOC stocks, which in the case of reduced tillage appeared to offset significant topsoil SOC gains. On the other hand, residue retention increased SOC stocks across the whole soil profile and may alleviate the neutral or negative effects of the other practices.

## 4.0 Discussion

### 4.1 Practices improve both yields and SOC stocks globally

Our global meta-analysis provides clear evidence that practices like reduced tillage, cover cropping, complex crop rotations and residue retention can increase topsoil SOC stocks and/or crop yields and frequently produce win-win outcomes (in 18.1% of observations). In contrast to many previous studies (Lobell et al., 2024; Ogle et al., 2019; Pittelkow et al., 2015), our results suggest that true trade-offs between SOC sequestration and crop yields (where one response is significantly positive while the other is significantly negative) are relatively rare (only 5.7% of observations). This suggests that many global and regional goals – such as the “4 per 1000” Initiative (Minasny et al., 2017), which aims to increase SOC stocks by 0.4% per year and stabilize or increase crop yields – may be achievable. However, it also highlights that win-wins are not guaranteed, meaning that much can be done to identify the optimal deployment of practices to maximize SOC sequestration and crop yields.

### 4.2 Heterogenous outcomes of practice implementation generate management trade-offs

While true trade-offs (where one outcome responds negatively) may be rare, the magnitude of topsoil SOC and yield responses depended on the type of practice implemented. Reduced tillage significantly increased SOC and had no effect on crop yields (Figure 2b). Within that, no-tillage increased SOC stocks more than minimum tillage, but minimum tillage increased crop yields while no-tillage did not (Figure 2e). The topsoil SOC effects were broadly consistent with previous studies (Ogle et al., 2019; Prairie et al., 2023; Shang et al., 2021; Sun et al., 2020); although we did not observe a negative effect of reduced tillage on yields as previous studies have (Pittelkow et al., 2015). Our results suggest that the decision over which method of reduced tillage to use generates a management trade-off: no-tillage may be optimal for soil carbon sequestration, but minimum tillage may be the optimal for farmers hoping to increase yields.

Within cropping system intensification practices, we found that cover cropping increased both topsoil SOC and crop yields while complex crop rotations only slightly increased topsoil SOC stocks (Figure 2c). The lack of literature consensus on the effects of cropping intensification (Abdalla et al., 2019; Lobell et al., 2024; McClelland et al., 2021; Poeplau & Don, 2015; Prairie et al., 2023; Zhao et al., 2022) may arise from the many complex decisions that practitioners must make when implementing cover cropping and crop rotations, such as cover crop functional group (Abdalla et al., 2019; King & Blesh, 2018). We found that using a grass cover crop produced a slightly higher SOC response than a legume cover crop. Conversely, using a legume cover crop produced a higher yield response (Figure 2c); however, the sample sizes and differences between mean values were small. Our findings contrast with a previous meta-analysis that showed legumes increased SOC stocks more than grasses (Jian et al., 2020). While Jian *et al* (2020) acknowledge the variable effects of cover crop functional group on SOC response, one explanation for this discrepancy could be that functional group inadequately captures key species-specific traits like biomass C:N ratio which may better predict SOC responses. Moreover, soil nitrogen availability and nitrogen application rate may influence the effectiveness of different cover crop species and functional groups. Our results again show a management trade-off where grass cover crops are preferable from the perspective of SOC sequestration, but legume cover crops maximize yield gains.

Lastly, residue retention, especially when incorporated into the soil, increased both topsoil SOC and crop yields (Figure 2b, f). Our results are consistent with the literature consensus about general residue retention (Liu et al., 2014; Pittelkow et al., 2015; Shang et al., 2021; Xia et al., 2018), suggesting that on a global scale residue retention is the individual practice that will most consistently produce win-wins whereas reduced tillage and intensification practices may be more likely to generate trade-offs.

### 4.3 Crop type and practice interactions generate further management trade-offs

Our results show that the crop type was an important determinant of the yield response, indicating that gains in maize and wheat yields are more common than rice and soybeans. Consequently, matching practice selection with crop type reveals further management trade-offs. For example, in both maize and wheat systems, residue retention led to the largest average topsoil SOC and yield gains of the individual practices (Table 1), as shown by Shang et al (2021). While in rice and soybean systems we found that residue retention led to the largest average SOC gains, intensification led to the largest average yield gains of the individual practices (Table 1). Thus, when choosing between individual practices, rice and soybean farmers may face a trade-off between maximizing yields (*via* intensification) or maximizing soil carbon sequestration (*via* residue retention).

**Table 1.**
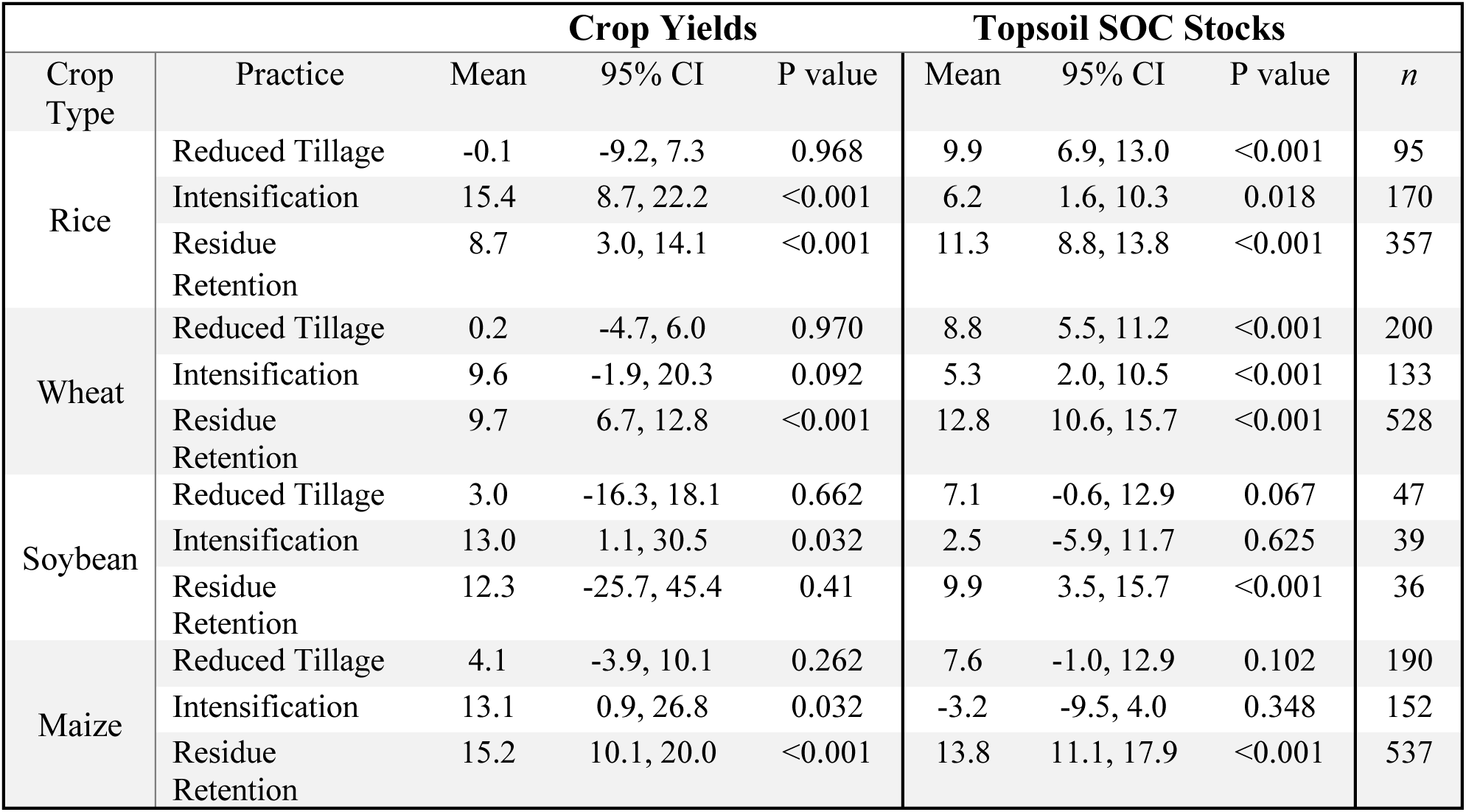
Effects of selected practices broken down by crop type. Mean values represent % change (converted from response ratio) of crop yields and topsoil SOC stocks for each combination of crop type and agricultural practice. 95% confidence intervals and p-values derived from 1000-iteration bootstrapping, and *n* reflects the sample size in each combination of crop and practice. Intensification = cover cropping and/or complex crop rotations. See Table S7 for full breakdown including practice stacking.

### 4.4 Practice stacking may alleviate management trade-offs associated with individual practices

Consistent with our expectations and the literature (McClelland et al., 2021; Pittelkow et al., 2015; Prairie et al., 2023), we found that combining multiple practices tends to have a beneficial effect on both topsoil SOC and yield outcomes. For example, combining reduced tillage or residue retention with intensification tended to increase the average yield response, and combining reduced tillage or intensification with residue retention helped overcome the negligible impacts of intensification and reduced tillage on SOC and yields, respectively. While previous meta-analyses have shown the benefits of practice stacking e.g., for improving the SOC benefit of cover cropping (McClelland et al., 2021; Pittelkow et al., 2015) or for improving the yield effect of reduced tillage (Pittelkow et al., 2015), few if any have assessed how the effects practice stacking vary by crop type and the extent to which stacking practices helps to maximize SOC and yield outcomes. By assessing the effects by crop type in our study, we found important management trade-offs (Table S7). For example, in both rice and maize systems, the yield gain is maximized by combining residue retention with intensification, whereas the average topsoil SOC gain is maximized by combining residue retention with reduced tillage. Thus, practice stacking is crucial not only to alleviating negative or neutral impacts of individual practices but also help to maximize crop-specific SOC and yield outcomes.

### 4.5 Environmental and agronomic conditions limit practice effectiveness to marginal lands

While practice choice and implementation, crop type, and their interaction are crucial determinants of topsoil SOC and yield outcomes, ultimately these decisions are constrained by climatic and soil conditions and farm management factors.

Climatic variables like temperature, aridity, and potential evapotranspiration were important predictors of the yield effects of all practices and of the topsoil SOC effects of reduced tillage (Table S8). Overall, we observed the greatest SOC response but the lowest yield response to all practices in arid regions (Figure 3a), meaning that arid regions are more likely to experience SOC gains and less likely to experience yield gains.

However, at the individual practice level, we find that reduced tillage performs best for both topsoil SOC and yields in more arid regions (Figure 4c) with low potential evapotranspiration (Figure 4b) and delivers win-wins in regions with an aridity index less than 0.5. Similarly, residue retention produces win-wins in areas with lower potential evapotranspiration less than 2000 (Figure 4e). Practices such as reduced tillage and residue retention may improve soil structure, water infiltration, and soil moisture retention, enabling crops to overcome water limitation, increasing both yields and carbon inputs (Ogle et al., 2019; Pittelkow et al., 2015; Xia et al., 2018). Given 45.1% of agricultural lands occupy areas with aridity index <0.5 and 89.1% of agricultural lands have a potential evapotranspiration less than 2000 (Figure S3), there is large spatial potential to implement sustainable practices in optimal conditions.

Similarly, all practices performed best in warmer climates (MAT > 20°C) (Figure 4a,d,g) likely because reduced tillage limits decomposition while residue and cover crop-derived carbon inputs compensate for faster organic matter turnover in warm conditions (Shang et al., 2021). However, cover crop effectiveness is limited in dry climates due to water competition with main crops (Bai et al., 2019; Lobell et al., 2024), explaining the lower overall yield responses in arid regions (Figure 3a). This suggests there may be limits to practice stacking, as combining other practices with cover cropping may provide no additional benefit or even be detrimental in arid climates.

We observed that the topsoil SOC change was negatively correlated with background (initial) SOC concentrations (Figure 3b), meaning greater gains were experienced in areas with low baseline carbon contents. The greatest SOC gains were found in soils with a baseline SOC content <0.6% (Figure 3c), which encompasses 76.4% of agricultural lands globally (Figure S3). Negative correlations between SOC change and initial SOC could be the result of statistical artifacts (Slessarev et al., 2023), giving us pause in speculating about the actual biogeochemical mechanism.

Interestingly, there was no such negative relationship between topsoil SOC response and the mineral-associated carbon saturation (Figure 3c). This indicates either that the soils are not MAOC-saturated (and thus can gain both POC and MAOC) or that the topsoil SOC gains are predominantly the result of POC gains rather than MAOC gains. Previous studies have found that sustainable agricultural practices have increased carbon storage in both POC and MAOC pools (Prairie et al., 2023). Thus, it is unclear (at least using global datasets) whether MAOC saturation can predict potential SOC changes at the global scale.

The SOC response remained constant across a gradient of nitrogen application rates, but the yield response was greatest in low-N systems (Figure 3d). This echoes previous results (Abdalla et al., 2019; King & Blesh, 2018; Zhao et al., 2022), suggesting that practices like cover cropping can help overcome nitrogen limitation (e.g., by growing N-fixing legumes) but this effect becomes masked when more externally-derived N is supplied. Interestingly, within cover cropping, crop type was an important predictor and influenced the nitrogen threshold at which the yield response became positive (Figure S5). Rice and maize showed positive yield responses when nitrogen application was below 160 kg N/ha while wheat had a lower threshold of 100 kg N/ha, and the soybean yield response is never significantly positive across the entire nitrogen gradient (though only becomes significantly negative when nitrogen application exceeds 160 kg N/ha). Most of harvested area globally exists in systems receiving less than these N application ‘thresholds’ (81.6% of rice, 76.3% of maize, and 62.9% of wheat harvested area) (Figure S4), suggesting broadly that N fertilizer usage may not inhibit the efficacy at the global scale.

### 4.6 Topsoil SOC changes are associated with increases in microbial biomass

Overall, sustainable agricultural practices increased microbial biomass carbon (MBC) (Figure 5a). These results are broadly consistent with the literature (Li et al., 2018; Zuber & Villamil, 2016) and likely reflect increased carbon inputs and soil microclimate conditions that favour microbial growth (Li et al., 2018; Liu et al., 2014; Prairie et al., 2023). However, the implications for SOC dynamics differ between practices: while residue retention and cropping intensification may increase topsoil SOC stocks partly through enhanced microbial activity that converts carbon inputs into soil organic matter (Liu et al., 2014; McClelland et al., 2021), reduced tillage may increase topsoil SOC primarily by reducing decomposition and maintaining soil structure (Shang et al., 2021), suggesting the concurrent increase in MBC could potentially constrain rather than accelerate topsoil SOC sequestration when reducing tillage.

### 4.7 Subsoil SOC responses are highly variable and may offset topsoil SOC gains

While subsoil carbon effects were highly variable (Figure 6), they adjusted our understanding of practice effectiveness beyond simply reducing statistical confidence. For reduced tillage, the inclusion of subsoil data transformed what appeared to be a significantly positive topsoil effect into a non-significant whole-profile response, indicating that subsoil losses likely offset topsoil gains. Similarly, cropping intensification showed no significant effect on SOC stocks across the full soil profile. This is consistent with previous research showing neutral or negative effects of some sustainable practices on subsoil carbon stocks (e.g., Ogle et al., 2019; Prairie et al., 2023). In contrast, residue retention maintained significant positive effects when subsoil responses were included, with significant subsoil gains bolstering topsoil increases to produce substantial carbon sequestration (+20.5%) across the whole profile. Thus, while residue retention consistently increases SOC stocks regardless of depth, subsoil responses may offset topsoil gains resulting from other practices such as reduced tillage.

**Figure 6.**
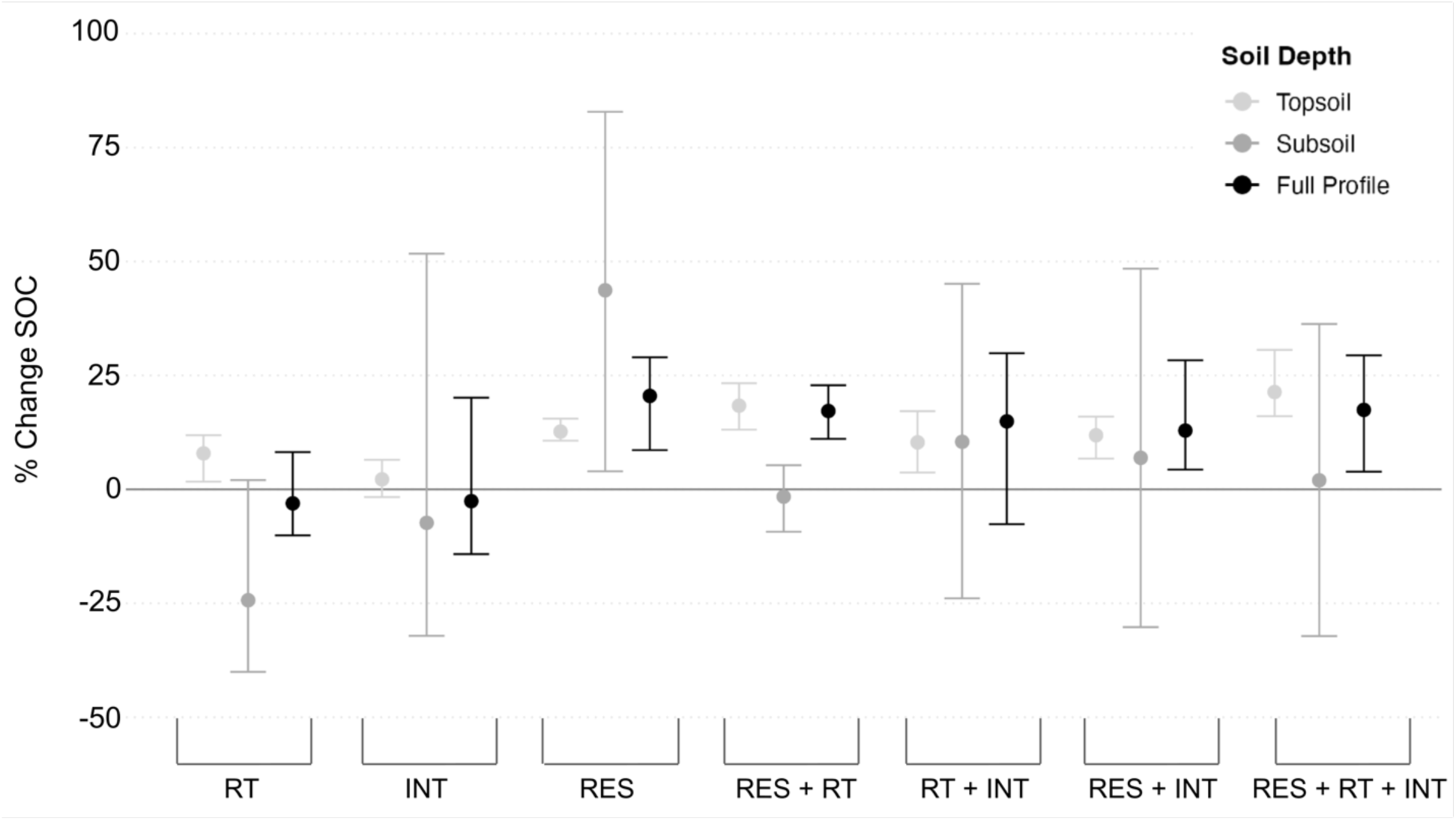
Soil carbon response by management practice and soil profile. Points represent mean estimates and error bars represent bootstrapped 95% confidence intervals, such that if the bars do not overlap the x-axis then the effects are significant. Topsoil, subsoil and full profile sample sizes are: Reduced tillage (RT: 532, 155, 158), Intensification (cover cropping and/or complex crop rotations) (INT: 494, 41, 44), Residue Retention (RES: 1458, 227, 243), Residue Retention and Reduced Tillage (RES + RT: 246, 43, 41), Reduced Tillage and Intensification (RT + INT: 69, 9, 9), Residue and Intensification (RES + INT: 131, 5, 17), all three practices (RES + RT + INT: 18, 6, 6).

### 4.8 Uncertainties in net emissions limit accurate estimates of climate benefits

Beyond subsoil responses, the effects of these practices on other on-farm and downstream greenhouse gas emissions are important uncertainties. For example, Qiu et al. (2024) found that cover cropping increased yields by 2.33% and SOC stocks by 6.46% but also increased N_2_O and CH_4_ emissions by 29.5% and 42.3%, respectively. Similarly, there is evidence that residue retention may increase CH_4_ emissions in rice paddies and N_2_O emissions in upland farms (Shang et al., 2021). Paustian et al. (2016) suggested that increases in microbial biomass carbon (and microbial activity) like those we observed (Figure 5) may drive increased CH_4_ and N_2_O emissions. However, sustainable practices could also reduce on-farm emissions by reducing fuel emissions (in the case of reduced tillage) or fertilizer-related emissions (by growing N-fixing cover crops or recycling nutrients through residue retention). As such, the net effect of these practices on climate change mitigation remains uncertain and is of vital importance for future research to enable effective implementation at the global scale.

## Conclusions

Our meta-analysis showed that, on a global scale, sustainable agricultural practices generally increased topsoil SOC stocks and crop yields, though the magnitude and significance of the change depended on the type of practice, crop, and the practice implementation method. By considering the interactions between crop types and multiple practices thought to bolster SOC, we reveal that management trade-offs primarily arise from determining which practice in which crop will maximize SOC and yield outcomes. For example, in rice systems, residue retention maximized topsoil SOC gains while cropping intensification maximized yield gains, though such trade-offs may be alleviated by combining multiple practices. Across the practices, win-wins were more likely in water- and nitrogen-limited systems over longer timeframes, indicating there is substantial global potential for deployment. However, uncertainties around subsoil SOC responses as well as on-farm and downstream emissions must be addressed. While we assessed relative yield and SOC changes, future work should determine the spatially explicit absolute changes in soil carbon stocks, total food production, knock-on emissions, and economic implications of sustainable agricultural practices to ensure effective policy design and large-scale implementation of sustainable practices.

## Acknowledgments

We thank all authors of the manuscripts that formed the basis of our meta-analysis. Juliana Kohli, Johanna Schoenecker, Martin Baur, Courtney Currier, and Sarayu Manoj also provided input on the manuscript and data analyses. Funding *via* UKRI grant EP/X042863/1 to AP.

## Author contributions

DE and AP designed the study and wrote the paper. DE conducted all data collection and analyses. PS and RP provided input on framing, interpretation of results and writing.

## Data availability statement

Data were collected from published materials that are freely accessible. Please see Table S2 of sources spatial datasets used in our study. The original, unedited dataset as well as other data subsets (e.g., subsoil and partial regression data frames) will be made available along with all the supporting code on a public repository (Zenodo) following acceptance.

