## Supplementary Information for "Identifying win-win opportunities and trade-offs for sustainable agriculture to improve agricultural productivity and soil carbon sequestration: A global meta-analysis"

### 1.0 - Methods

#### 1.1 - Literature Search

We comprehensively searched Web of Science and SCOPUS for studies that quantified both the yield and soil carbon impacts of four agricultural practices: cover cropping, complex crop rotations, reduced tillage, and crop residue retention (Table S1). We searched for studies that investigated the impact of any of these practices on soil carbon concentrations or stocks *and* yields of four main staple crops (maize, wheat, rice, and soybeans) (Lists S1 & S2). The initial searches included other crops such as potatoes, sugar cane, tomatoes, and cotton, but the final analysis focused on the four main staple crops.

**Table S1 – Definitions of agricultural practices assessed in the meta-analysis**

| **Agricultural Practice** | **Definition** |
|---|---|
| Cover cropping | "the planting of non-cash crops between the primary growing seasons" (Lobell et al., 2024) |
| Complex crop rotations | Altering the sequence of crop species growing in subsequent growing seasons. Typically involves either a) increasing number of cropping seasons per year, or b) increasing the number of crop species in a rotation. |
| Reduced tillage | Reducing soil disturbance. Involves converting from conventional tillage (CT; typically mechanical inversion) to minimum tillage (MT; reduced disturbance) or no-tillage (NT; no disturbance). |
| Improved residue management (residue retention) | Retaining or incorporating crop residues (instead of burning or removing them). Generally, refers to the use of on-farm residues, but also includes application of externally derived, off-farm residues. |

**List S1 - Search string used to gather articles from Web of Science**

*TS=(wheat OR maize OR corn OR rice OR soy* OR potato* OR sugar* OR cotton OR tomato*) AND TS=("minimum till*" OR "reduced till*" OR "surface till*" OR "no till*" OR "conservation till*" OR "conservation agriculture" OR no-till* OR min-till* OR reduced-till* OR surface-till* OR conservation-till* OR "cover crop" OR "green manure*" OR interplant* OR "companion crop*" OR "crop rotation*" OR "catch crop*" OR mulch* OR residue* OR stubble OR stalk* OR stover* OR straw) AND TS=(carbon OR SOC OR "organic matter") AND TS=(yield* OR production)*

**List S2 – Search string used to gather articles from SCOPUS**

*TITLE-ABS-KEY (wheat OR maize OR corn OR rice OR soy* OR potato* OR sugar* OR cotton OR tomato* ) AND TITLE-ABS-KEY ( "minimum till*" OR "reduced till*" OR "surface till*" OR "no till*" OR "conservation till*" OR "conservation agriculture" OR no-till* OR min-till* OR reduced-till* OR surface-till* OR conservation-till* OR "cover crop" OR "green manure*" OR interplant* OR "companion crop*" OR "crop rotation*" OR "catch crop*" OR mulch* OR residue* OR stubble OR stalk* OR stover* OR straw ) AND TITLE-ABS-KEY ( carbon OR soc OR "organic matter*" ) AND TITLE-ABS-KEY ( yield* OR production)*

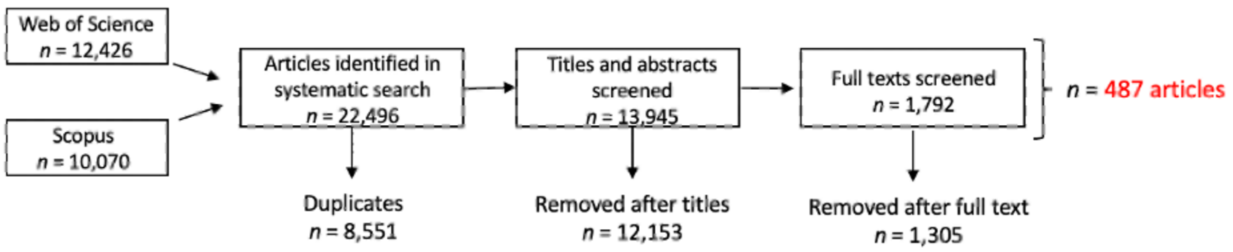


**Figure S1 – PRISMA flow diagram of paper screening.**

<u>1.2 - Study Selection Criteria</u>

We screened publications following the Preferred Reporting Items for Systematic Reviews and Meta-Analyses (PRISMA) guidelines with the following inclusion criteria (Moher et al., 2009). The study was conducted in the field with at least one full field season of treatment and control monitoring. The study reported both crop yield data (average or final year yields) for one of the four target crops **as well as** soil carbon data that can be converted to carbon stocks (such as %, g/kg, or tons/ha). The study compared a single sustainable practice (e.g., reduced tillage) or combination of practices (e.g., reduced tillage plus cover cropping) to the lack of that practice (control). Initial climate and soil parameters must have been the same in the control and treatment plots and the measurements in each plot must have been made at the same spatial/temporal scales. The study must have provided location data that can be converted to latitude and longitude. If multiple publications reported results from the same study site (e.g., at different times in a long-term experiment, or different trials at the same site), we coded them as the same study. The study must have specified the type of crop grown, the soil sampling depth, and the length of the experimental treatments. If the study reported different crops from the same treatments (e.g., reported yields for both rice and wheat in a rice-wheat rotation), we included each crop independently (Zhao et al., 2022).

### 1.3 Data Compilation and Management

#### *1.3.1 - Data Collection*

For each study, we collected the means, number of replications (sample size) and estimates of variability (standard deviation, standard error, coefficient of variation) for the SOC and yield parameters in control (conventional) and treatment (regenerative) groups. When data were only available in figures, we used WebPlotDigitizer (Rohatgi, 2024) to extract the data. If a measure of variability was reported (e.g., error bars) but the exact measure was not specified, it was not included in the data. Nitrogen application rate was standardised to kg N/ha using the International Fertilizer Association conversion rates (https://www.ifastat.org/converter/fertilizer-converter/).

We primarily used data on final year yields (e.g., a 6-year study that reported yields in the 6$^{th}$ year) as they were more commonly reported than mean yields. When mean yields were the only measurement provided, they were used in place of final year yields. Some North American yields were reported in bushels per acre, which was converted to tonnes/ha per Ulery & Drees (2008).

#### *1.3.2 - Bulk density interpolation*

Prior to aggregation, missing bulk density values in the original dataset were filled using treatment-specific means calculated from measured data. This was to preserve the effects of the practices on bulk density rather than using a single imputed value for both control and treatment groups. Indeed, we found the average bulk density in the control observations (1.36) to be slightly higher than in the treatment observations (1.34). These values were then used to fill in missing bulk density values prior to carbon stock calculation.

*1.3.3 - Depth Aggregation*

We developed a comprehensive depth aggregation framework to consolidate soil measurements from multiple depth intervals into standardized topsoil (~0-30 cm) and subsoil (>30 cm) layers while preserving the physical and chemical integrity of soil properties through appropriate mathematical treatments. We classified soil depth intervals into topsoil and subsoil categories using a 30 cm threshold as the primary delineation, with a 5 cm overlap tolerance (Prairie et al., 2023), allowing depth ranges like 0-35 cm or 15-35 cm to be classified as topsoil, and 25-40 cm as subsoil. We also applied a "75% rule" whereby depth intervals where ≥75% of the range fell within one layer were assigned to that layer (e.g., 0-40 cm → topsoil). The "full profile" was defined as any depth-aggregated observation where the maximum depth exceeded 30 cm (i.e., where the soil sampling included both topsoil and subsoil increments).

For the aggregation process specifically, we used bulk density values from the Harmonized World Soil Database (HWSD) (FAO & IIASA, 2023) based on layer classification to calculate soil mass for mass-weighted averaging. We employed different mathematical approaches depending on the nature of each soil property. Soil organic carbon stocks were summed within each layer as additive properties, while standard deviations were combined using root sum of squares for proper error propagation ($Combined\_SD = \sqrt{(\Sigma(SD\_i^2))}$). For intensive properties like pH, clay content, and nutrient concentrations, we calculated mass-weighted means where $Layer\_Mass\_i = BD\_i \times (Lower\_Depth\_i - Upper\_Depth\_i) \times 1000$, and $Weighted\_Mean = \Sigma(Property\_i \times Layer\_Mass\_i) / \Sigma(Layer\_Mass\_i)$. This approach ensures that thicker or denser soil layers contribute proportionally more to the aggregated value.

For subsoil and full soil profile observations, the depth aggregation approach required additional considerations given that soil bulk density generally increases with depth (Panagos et al., 2024). Prior to aggregation, missing bulk density values were filled using layer-specific means calculated from measured data: control topsoil mean = 1.34 g $cm^{-3}$, control subsoil mean = 1.50 g $cm^{-3}$, treatment topsoil mean = 1.32 g $cm^{-3}$, treatment subsoil mean = 1.47 g $cm^{-3}$. This approach preserved both treatment effects and the natural increase in bulk density with depth. For the aggregation process, subsoil observations used subsoil-specific bulk density values from the HWSD for mass calculations and gap-filling. Full soil profile observations were defined as any depth-aggregated comparison where the maximum sampling depth exceeded 30 cm, indicating that both topsoil and subsoil increments were included in the soil sampling. For these profiles, separate aggregated values were calculated for each layer (topsoil and subsoil) within the same comparison. The mass-weighted aggregation was particularly important for full profiles, as it properly accounted for the typically higher bulk density and different physicochemical properties of deeper soil layers when calculating profile-integrated values for soil organic carbon stocks and other properties.

For each aggregated layer, we calculated minimum depth (shallowest sampling point), maximum depth (deepest sampling point), and mean depth (midpoint of the entire layer range). The aggregation process resulted in separate rows for topsoil and subsoil within each comparison, with aggregated SOC stocks, mass-weighted soil properties, and depth summaries. Non-depth-varying metadata (e.g., study location, treatment type) were preserved from the first measurement within each layer.

1.4 - Spatial Data

Missing data for environmental covariates (e.g., soil pH, clay content, aridity index) were extracted from global gridded datasets using the latitude and longitude provided by or estimated from the studies. Sources for these global datasets are in Table S2. The Climate Classification layer was created using data from the UN FAO Global Agro-Ecological Zones (GAEZ) (Ulery & Drees, 2008). Specifically, historical (1981-2010) data regarding thermal and moisture regimes were combined following a similar approach to that outlined by Smith et al. (2007) to standardise the thermal and moisture regimes into climatic categories. We aggregated the thermal regime into four categories: Tropical (TRC1 & TRC2), Subtropical (TRC3, TRC4, & TRC5), Temperate (TRC6 & TRC7), and Boreal (TRC8 & TRC9). We excluded arctic environments (TRC10) as agriculture is exceedingly rare in these environments. We similarly aggregated the moisture regime into two categories: Dry and Moist, where the length of the growing period (LGP) is <180 days or >180 days, respectively. These two reclassified thermal and moisture regime datasets were combined to generate one map with eight combinations: Tropical Dry, Tropical Moist, Subtropical Dry, Subtropical Moist, Temperate Dry, Temperate Moist, Boreal Dry, and Boreal Moist.

The "yield gap" factor was derived from the FAO GAEZ data on crop yield achievement ratios, which represents the gap between actual yields and potential yields. Thus, a higher yield achievement ratio indicates a lower yield gap, and vice versa. The GAEZ data provides yield achievement ratios for irrigated and rainfed areas across multiple crop categories: all crops combined, cereals, and our four staple crops (maize, wheat, soybean, and wetland rice). For each sampling location, yield gap values were extracted using geographic coordinates and categorized into seven classes ranging from <10% to >85% achievement ratio. The assignment of yield gap values followed a conditional logic based on irrigation status: locations with irrigation were assigned irrigated yield gap values, while non-irrigated or unknown irrigation status locations received rainfed yield gap values. For crop-specific analyses, an additional conditional layer was applied where the appropriate crop-specific yield gap (maize, wheat, soybean, or rice) was assigned based on the primary crop type at each location, with wheat categories consolidated to include durum wheat, bread wheat, spring wheat, and winter wheat varieties, and rice categories including basmati as well as both early and late season rice varieties.

MAOC Saturation is derived from Georgiou et al. (2022) and refers to the realized amount of mineral associated organic carbon (MOC) divided by the potential amount of mineral-associated carbon ($MOC_{max}$).

**Table S2 – Sources of environmental and agronomic covariates**

| Covariate | Source | Resolution | Modifications |
|---|---|---|---|
| Mean annual temperature (MAT) | WorldClim 2.0<br>(Fick & Hijmans, 2017) | 30" | Averaged the mean monthly temperatures |
| Minimum temperature (Min T) | WorldClim 2.0 | 30" | Averaged the mean monthly minimum temperatures |
| Maximum temperature (Max T) | WorldClim 2.0 | 30" | Averaged the mean monthly maximum temperatures |
| Mean annual precipitation (MAP) | WorldClim 2.0 | 30" | Summed the mean monthly precipitation values |
| Mean monthly precipitation (MMP) | WorldClim 2.0 | 30" | Averaged the mean monthly precipitation values |
| Elevation | WorldClim | 30" | |
| Aridity index | (Zomer et al., 2022) | 30" | |
| Potential evapotranspiration (PET) | (Zomer et al., 2022) | 30" | |
| Climate class | GAEZ<br>(Fischer et al., 2021) | 5' | See above |
| Median slope | GAEZ | 5' | Excluded "Water". Combined >45% and 30-45% into >30%, and 0-0.5% and 0.5-2% into <2%. |
| pH | HWSD<br>(FAO & IIASA, 2023) | 30" | |
| Clay content | HWSD | 30" | |
| Silt content | HWSD | 30" | |
| Sand content | HWSD | 30" | |
| Bulk density | HWSD | 30" | |
| Cation exchange capacity in the clay fraction (CEC) | HWSD | 30" | |
| Background SOC | HWSD | 30" | |
| Drainage class | HWSD | 30" | Aggregated 7 classes to create 5. Combined "Very Poor" and "Poor" as well as "Excessive" and "Somewhat Excessive". |

| | | | |
|---|---|---|---|
| Soil texture | HWSD | 30" | Removed class 0 (None). 3 classes: Fine, Medium, Coarse |
| Yield gap | GAEZ | 5’ | Aggregated <10% and 10-25% into <25%. Aggregated >85% and 70-85% into >70%. |
| MAOC Saturation | (Georgiou et al., 2022) | 0.5 degrees | Divided MOC (realized mineral associated organic carbon) by $MOC_{max}$ (potential mineral associated organic carbon), such that a saturatinvalue of 1 means that all of the potential mineral-associated carbon storage has been achieved. |

### 1.5 - Meta-Analysis

#### *1.5.1 - Practice-specific subsets*

While some of the analyses used the full paired topsoil and yield dataset (n = 2975) (Figures 1, 2a, 2b, and 3; Table 1), some analyses used practice-specific subsets that focused on specific practices (Figures 2c-f, 4). In these cases, the practice-specific subsets included all observations where that practice is adopted, regardless of whether other practices are also adopted. In other words, the practice-specific subsets were not limited to observations where the only difference between control and treatment is the one practice. So, for example, an observation that compared a control with conventional tillage and residue removal to a treatment with no-tillage and residue retention (which would be an example of practice stacking as both reduced tillage and residue retention are used) would be included in both the “Reduced Tillage” and “Residue Retention” subsets. Outlier removal (e.g., yield and SOC response ratio outliers) was performed at the level of the practice-specific subsets.

#### *1.5.2 - Variable Importance – Discussion of Metaforest Cross-Validation methods*

We calculated both the out-of-bootstrap $R^2$ ($R^2_{oob}$) and the tenfold cross-validated $R^2$ ($R^2_{cv}$). The model hyperparmeters, $R^2_{oob}$, and $R^2_{cv}$ for the top trained models are shown in Table S3. The $R^2_{oob}$ and $R^2_{cv}$ ranged from 0.10 to 0.43 and 0.14 to 0.48, respectively, indicating that the top models had reasonable predictive performance.

When using random forest machine learning approaches like *metaforest* (Lissa, 2020), the cross-validation method used has implications for hyperparameter tuning and performance estimation. The two main approaches applied to meta-analytic datasets are “traditional” 10-fold cross validation in which observations are randomly assigned to folds, and “clustered” cross-validation which groups all observations from the same study site so that they can't appear in both the training and test sets. In a meta-analytic context with large random effects and study-site clustering, traditional cross-validation may lead to models that are over-fit on site-specific factors since observations from the same study site can be used in both the training and test sets, leading to an overestimation of model performance. Clustered cross-validation better captures this non-independence of observations and may lead to a more conservative estimate of model performance. Despite this, numerous previous meta-analyses have used traditional cross-validation with *metaforest* despite presumably having study site clustering of data (Terrer et al., 2021; Zhao et al., 2022).

We performed both traditional and clustered cross-validation to determine how the cross-validation method affected hyperparameter tuning, model performance, and variable importance (Table S3). The cross-validated $R^2$ was consistently lower in the traditionally cross-validated models compared to the clustered cross-validated models, indicating that traditional cross-validation may overestimate model performance by ignoring study site data clustering and strong random effects. Nevertheless, using a threshold of 0.7 (70/100) to identify important predictors, both cross validation methods identified the same important predictors. This suggests that although our assessment of model performance is strongly influenced by cross-validation method, the "important" predictors are consistently important regardless of cross-validation method.

**Table S3 – Comparison of traditional and clustered cross-validation methods**. Shows the hyperparameters (weights, number of candidate variables considered at each split [mtry], and minimum node size) of the top model, as well as the cross-validated and out-of-bootstrap $R^2$ values for both the top models derived from “traditional” 10-fold cross-validation and “clustered” cross-validation methods.

| | **Traditional cross-validation** | | | **Clustered cross-validation** | | |
|---|---|---|---|---|---|---|
| Submodel | Hyperparameters | $R^2_{cv}$ | $R^2_{oob}$ | Hyperparameters | $R^2_{cv}$ | $R^2_{oob}$ |
| Cover Crop Yields | Weights: Random<br>Minimum node size: 2<br>Mtry: 2 | 0.48 | 0.43 | Weights: Fixed<br>Minimum node size: 3<br>Mtry: 2 | 0.31 | 0.42 |
| Cover Crop SOC | Weights: Uniform<br>Minimum node size: 2<br>Mtry: 1 | 0.24 | 0.19 | Weights: Uniform<br>Minimum node size: 6<br>Mtry: 1 | 0.17 | 0.19 |
| Rotation Yields | Weights: Uniform<br>Minimum node size: 2<br>Mtry: 2 | 0.36 | 0.28 | Weights: Uniform<br>Minimum node size: 2<br>Mtry: 1 | 0.30 | 0.27 |
| Rotation SOC | Weights: Random<br>Minimum node size: 5<br>Mtry: 2 | 0.14 | 0.10 | Weights: Random<br>Minimum node size: 2<br>Mtry: 1 | 0.15 | 0.10 |
| Tillage Yields | Weights: Fixed<br>Minimum node size: 4<br>Mtry: 2 | 0.18 | 0.14 | Weights: Fixed<br>Minimum node size: 5<br>Mtry: 2 | 0.06 | 0.14 |
| Tillage SOC | Weights: Uniform<br>Minimum node size: 4<br>Mtry: 3 | 0.39 | 0.40 | Weights: Uniform<br>Minimum node size: 5<br>Mtry: 2 | 0.10 | 0.40 |
| Residue Yields | Weights: Uniform<br>Minimum node size: 2<br>Mtry:2 | 0.38 | 0.39 | Weights: Fixed<br>Minimum node size: 6<br>Mtry: 2 | 0.06 | 0.33 |
| Residue SOC | Weights: Uniform<br>Minimum node size: 5<br>Mtry: 4 | 0.39 | 0.39 | Weights: Fixed<br>Minimum node size: 5<br>Mtry:2 | 0.08 | 0.15 |

#### 1.6 - Model Assumptions

For all univariate mixed effects (*rma.mv*) models and multivariate partial regressions (*rma* models), we used statistical tests to assess publication bias and other model assumptions.

Egger’s regression (Egger et al., 1997) suggested that many of the funnel plots were skewed (and therefore there was significant publication bias), but in most cases the trim-and-fill method did not add any observations. In the few cases where it did, the trim-and-fill method illustrated that any publication bias was producing an underestimation. Furthermore, the fail-safe numbers (Rosenberg, 2005) for all datasets were sufficiently large (range: 27,618 to 29,934,282) and we thus concluded that the models were robust to publication bias (Table S4). The discrepancy between Egger's test and the trim-and-fill or Fail-Safe N Analysis may arise because Egger's test struggles to distinguish between true publication bias and the extreme heterogeneity in our data ($I^2$ values > 95%) (Nakagawa et al., 2022).

Our analysis suggested that many of the univariate mixed effects models used to generate forest plots and multivariate models used for partial regressions violated certain assumptions, particularly the assumption of residuals being normally distributed (Tables S5 & S6). To resolve this, we bootstrapped the fitted coefficients with 1000 iterations, sampling with replacement by study site. Point estimates and error bars in the forest plots (and lines and error bars in regressions) represent the mean estimate of the 1000 iterations and the 95% confidence intervals based on the distribution of the 1000 estimates. Yield and soil carbon responses were considered significant if the 95% confidence interval did not overlap with zero.

**Table S4 – Publication Bias Tests.** Table shows the sample size of each subset used to assess publication bias. See Section 1.5.1 for details of practice-specific subsets. $I^2$ indicates the heterogeneity of the datasets. Egger's regression test used to statistically assess for publication bias – if $p < 0.05$ then publication bias is considered significant. Fail-Safe N refers to the number of null observations that would need to be added to reduce the statistically significant meta-analytic effect size to non-significance, and the threshold value is 5n + 10 (Fragkos et al., 2014; Zhao et al., 2022).

| Submodel | Sample size (n) | Overall effect [95% CI] | $I^2$ (%) | Egger's p | Trim-Fill Missing Observations | Trim-fill adjusted effect | Fail-Safe N |
|---|---|---|---|---|---|---|---|
| Full Yield | 2975 | 0.091 [0.083, 0.099] | 96.85% | <0.0001 | 0 | NA | 9,400,035 |
| Full SOC | 2975 | 0.091 [0.086, 0.096] | 98.94% | <0.0001 | 0 | NA | 29,934,282 |
| Cover Crop Yield | 400 | 0.172 [0.144, 0.200] | 96.83% | 0.7059 | 95 (right side) | 0.256 | 456,051 |
| Cover Crop SOC | 400 | 0.097 [0.082, 0.111] | 92.23% | 0.0031 | 0 | NA | 234,351 |
| Rotation Yield | 334 | 0.139 [0.107, 0.171] | 96.55% | 0.5021 | 0 | NA | 186,611 |
| Rotation SOC | 334 | 0.047 [0.032, 0.062] | 99.34% | 0.0028 | 0 | NA | 51,381 |
| Tillage Yield | 862 | 0.0229 [0.009, 0.037] | 96.87% | 0.2292 | 0 | NA | 97,569 |
| Tillage SOC | 861 | 0.089 [0.079, 0.098] | 99.26% | 0.0371 | 0 | NA | 3,341,757 |
| Residue Yield | 1842 | 0.089 [0.079, 0.098] | 99.26% | 0.004 | 0 | NA | 5,787,208 |
| Residue SOC | 1841 | 0.110 [0.104, 0.117] | 98.75% | <0.001 | 0 | NA | 16,881,580 |
| MBC Full | 326 | 0.189 [0.160, 0.219] | 97.62% | <0.001 | 90 (right side) | 0.276 | 592,569 |
| MBC Tillage | 66 | 0.236 [0.162, 0.309] | 98.36% | 0.086 | 14 (right side) | 0.332 | 45,440 |
| MBC Residue | 208 | 0.184 [0.155, 0.212] | 96.19% | <0.001 | 39 | 0.238 | 234,671 |
| Subsoil SOC | 498 | 0.022 [0.008, 0.036] | 98.69% | 0.309 | 0 | NA | 27,618 |
| Full profile SOC | 531 | 0.077 [0.069, 0.085] | 98.01% | 0.003 | 0 | NA | 1,298,102 |

**Table S5 – Model assumption testing for univariate mixed-effects models.** Univariate models used for forest plots (Figures 2,3,5 and 6 in the main text). See Section 1.5.1 for details of practice-specific subsets. Distribution of residuals assessed using Q-Q plots (not shown) and Shapiro-Wilk test ($p < 0.05$ indicates that residuals are not normally distributed). Homoscedasticity (constant variance) was tested with residuals vs fitted plots (not shown) and the variance ratio (the ratio of the variances in the residual quartiles with the maximum and minimum variances). A threshold of 3 is used here with the residuals vs fitted plots to determine if the variance is constant (variance ratio < 3) or if there is heteroscedasticity (Hartley, 1950).

| Model | Data | Sample Size | Shapiro p-value | Variance Ratio | Normality Violated | Homoscedasticity Violated |
|---|---|---|---|---|---|---|
| Aridity Yield | Full | 2975 | < 0.001 | 1.08 | YES | NO |
| Aridity SOC | Full | 2975 | <0.001 | 1.16 | YES | NO |
| Background SOC Yield | Full | 2934 | <0.001 | 2.00 | YES | NO |
| Background SOC SOC | Full | 2934 | <0.001 | 1.22 | YES | NO |
| MAOC Saturation Yield | Full | 2814 | <0.001 | 1.46 | YES | NO |
| MAOC Saturation SOC | Full | 2814 | <0.001 | 1.05 | YES | NO |
| Nitrogen rate Yield | Full | 2971 | <0.001 | 1.79 | YES | NO |
| Nitrogen rate SOC | Full | 2971 | <0.001 | 1.46 | YES | NO |
| Study duration Yield | Full | 2975 | <0.001 | 1.22 | YES | NO |
| Study duration SOC | Full | 2975 | <0.001 | 1.12 | YES | NO |
| Crop type Yield | Full | 2975 | <0.001 | 2.91 | YES | NO |
| Crop type SOC | Full | 2975 | <0.001 | 2.81 | YES | NO |
| Management Yield | Full | 2948 | <0.001 | 1.27 | YES | NO |
| Management SOC | Full | 2948 | <0.001 | 1.38 | YES | NO |
| Crop x Management Yield | Full | 2948 | <0.001 | 2.03 | YES | NO |
| Crop x Management SOC | Full | 2948 | <0.001 | 2.36 | YES | NO |
| Intensification Type Yield | Full | 479 | <0.001 | 1.33 | YES | NO |
| Intensification Type SOC | Full | 479 | <0.001 | 1.04 | YES | NO |
| CC Functional Yield | Cover crop | 400 | <0.001 | 1.75 | YES | NO |
| CC Functional SOC | Cover crop | 400 | <0.001 | 1.85 | YES | NO |
| Control Rotation Yield | Rotation | 334 | <0.001 | 3.22 | YES | YES |
| Control Rotation SOC | Rotation | 334 | 0.0024 | 2.76 | YES | NO |

| | | | | | | |
|---|---|---|---|---|---|---|
| Tillage type Yield | Tillage | 861 | <0.001 | 1.60 | YES | NO |
| Tillage type SOC | Tillage | 861 | <0.001 | 1.30 | YES | NO |
| Residue Location Yield | Residue | 1606 | <0.001 | 1.50 | YES | NO |
| Residue Location SOC | Residue | 1606 | <0.001 | 1.14 | YES | NO |
| MBC Management | MBC Full | 324 | <0.001 | 2.75 | YES | YES |
| MBC Tillage Type | MBC Tillage | 65 | 0.027 | 1.97 | YES | NO |
| MBC Residue Location | MBC Residue | 201 | <0.001 | 1.64 | YES | NO |
| MBC Aridity | MBC Full | 322 | <0.001 | 1.71 | YES | NO |
| Pure subsoil Management | Pure subsoil | 486 | <0.001 | 3.52 | YES | YES |
| Full profile Management | Full profile | 518 | <0.001 | 2.58 | YES | YES |

**Table S6 – Model assumption testing for multivariate fixed-effects models**. See description for Table S5, in this case the models are the partial regression models used for Figure 4. The variables included in each model are shown in Table S9.

| Model | Data | Sample Size | Shapiro p-value | Variance ratio | Normality violated | Homoscedasticity violated |
|---|---|---|---|---|---|---|
| Cover crop yields | Cover Crop | 368 | <0.001 | 8.14 | YES | YES |
| Cover crop SOC | Cover Crop | 368 | <0.001 | 5.67 | YES | YES |
| Rotation yields | Rotation | 285 | <0.001 | 4.34 | YES | YES |
| Rotation SOC | Rotation | 285 | <0.001 | 3.96 | YES | YES |
| Tillage yield | Tillage | 796 | <0.001 | 1.66 | YES | NO |
| Tillage SOC | Tillage | 796 | <0.001 | 3.73 | YES | YES |
| Residue yield | Residue | 1725 | <0.001 | 3.84 | YES | YES |
| Residue SOC | Residue | 1725 | <0.001 | 3.52 | YES | YES |

### 2.0 - Results

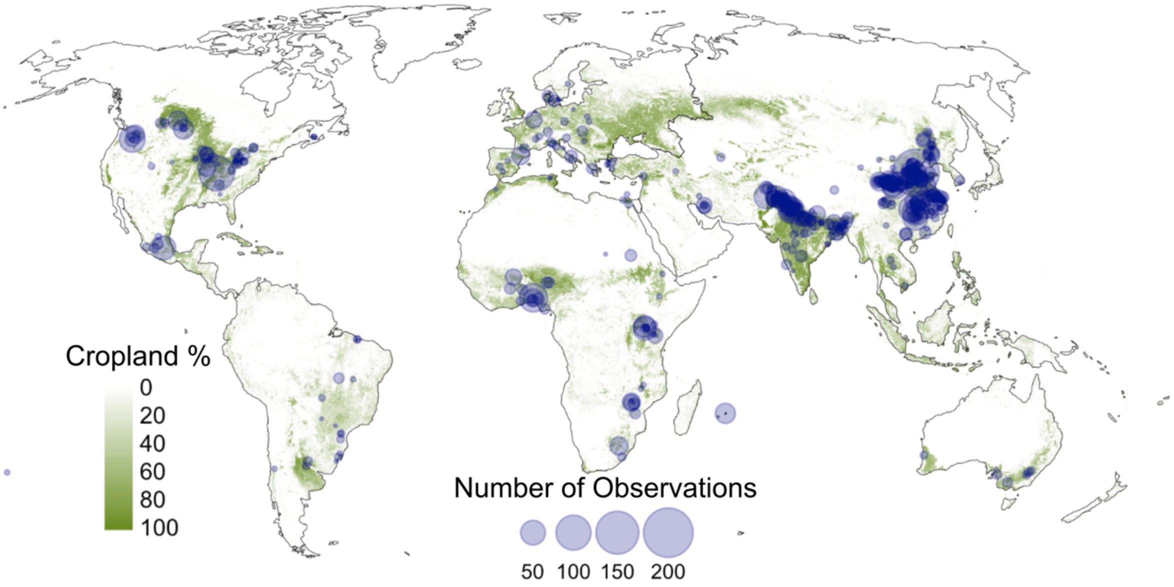


**Figure S2 - Map of depth-aggregated observations superimposed on global cropland.** Global cropland map derived from FAO Global Land Cover Share Database (Latham et al., 2014).

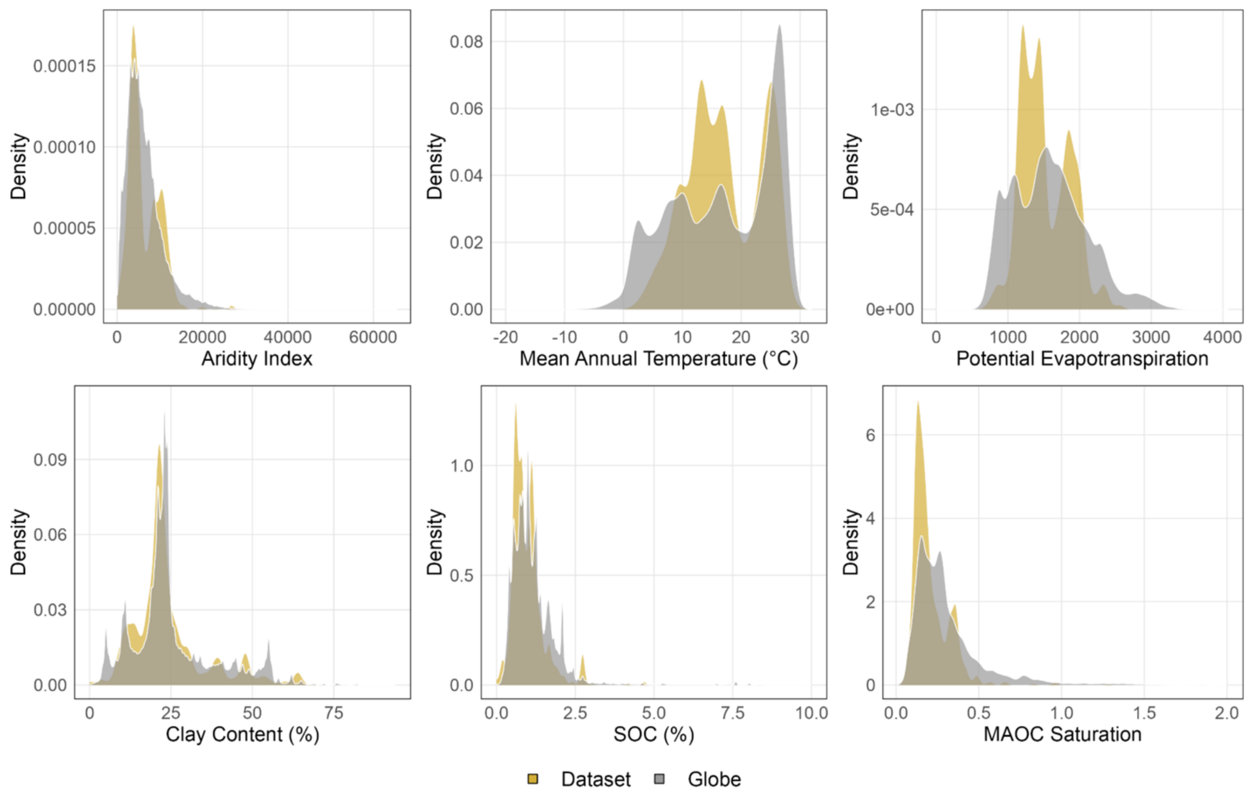


**Figure S3 – Representativeness of global dataset in terms of certain environmental covariates.** Density graphs of the distribution of variable values in our paired topsoil SOC and yield dataset (yellow) and the global values (grey), restricted to between –60 and +60 latitudes to exclude boreal, Arctic and Antarctic regions. Croplands defined as pixels with >1% agricultural coverage from by the FAO Global Land Cover Share Database (Latham et al., 2014) at 5 arc-minute resolution. The FAO Global Land Cover Share Database identifies cropland on a proportional basis, so the global distributions were weighted by the actual land cover (in hectares). Climate variables (top row) include aridity index multiplied by 10,000, potential evapotranspiration (Zomer et al., 2022), and mean annual temperature (°C) (Fick & Hijmans, 2017). Soil variables (bottom row) include clay content (%), soil organic carbon (%) (FAO & IIASA, 2023), and mineral-associated organic carbon (MAOC) saturation – defined as the realized amount of mineral-associated carbon relative to the potential amount of mineral-associated carbon (Georgiou et al., 2022).

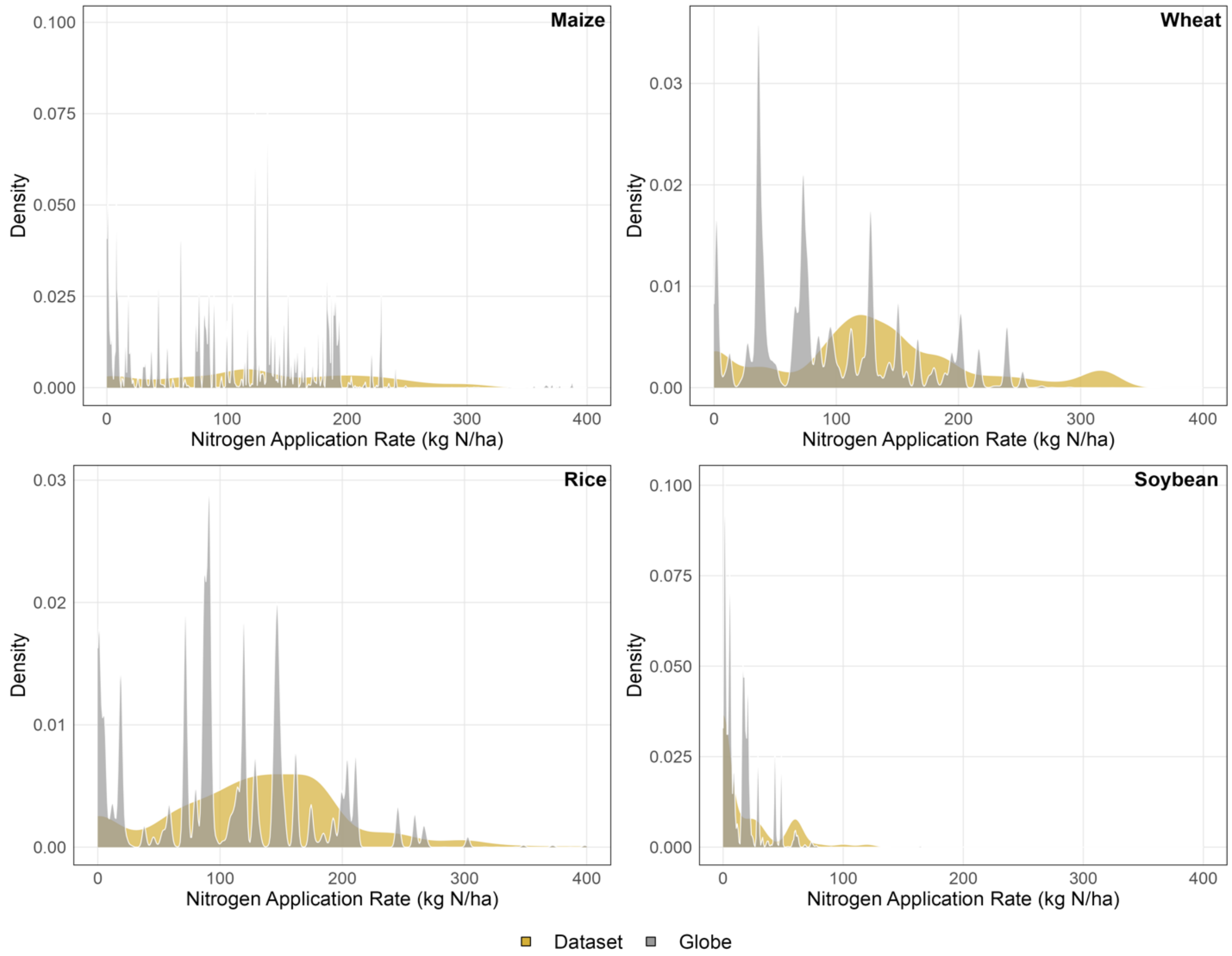


**Figure S4 – Nitrogen application rates by crop: comparison of dataset and global cropland areas**. Density distributions of nitrogen application rates comparing study sites (yellow) to global crop-specific areas (grey) for maize, wheat, rice, and soybean. Global distributions are weighted by harvested area (hectares) from CROPGRIDS (Tang et al., 2024). Harvested area differs from cropping area in that it double-counts areas in which the same crop is grown twice (or three times) in a year. Nitrogen application rates from NPKGRIDS (Nguyen et al., 2024) represent synthetic fertilizer application only (kg N/ha/year) at 0.05° resolution. Global data only include pixels where each specific crop is grown (harvested area > 0 hectares), while dataset observations are filtered by crop type.

Table S7 – **Effects of sustainable practices broken down by crop type**. Mean values represent % change (converted from response ratio) of crop yields and topsoil SOC stocks. 95% confidence intervals and p-values derived from 1000-iteration bootstrapping, and *n* reflects the sample size in each combination of crop and practice. RT = Reduced Tillage, RES = Residue Retention, INT = Cropping Intensification (cover cropping and/or complex crop rotations). "+" denotes the stacking of multiple practices.

| | | **Crop Yields** | | | **Topsoil SOC Stocks** | | | |
|---|---|---|---|---|---|---|---|---|
| Crop Type | Practice | Mean | 95% CI | P value | Mean | 95% CI | P value | *n* |
| Rice | RT | -0.1 | -9.2, 7.3 | 0.968 | 9.9 | 6.9, 13.0 | <0.001 | 95 |
| | INT | 15.4 | 8.7, 22.2 | <0.001 | 6.2 | 1.6, 10.3 | 0.018 | 170 |
| | RES | 8.7 | 3.0, 14.1 | <0.001 | 11.3 | 8.8, 13.8 | <0.001 | 357 |
| | RES + RT | 4.4 | -6.0, 14.8 | 0.45 | 18.7 | 14.0, 24.8 | <0.001 | 48 |
| | RT + INT | NA | NA | NA | NA | NA | NA | 0 |
| | RES + INT | 16.9 | 11.8, 22.4 | <0.001 | 11.7 | 6.1, 15.7 | <0.001 | 70 |
| | RES + RT + INT | 5.3 | -21.9, 17.8 | <0.001 | 20.3 | 16.8, 25.5 | <0.001 | 3 |
| Wheat | RT | 0.2 | -4.7, 6.0 | 0.970 | 8.8 | 5.5, 11.2 | <0.001 | 200 |
| | INT | 9.6 | -1.9, 20.3 | 0.092 | 5.3 | 2.0, 10.5 | <0.001 | 133 |
| | RES | 9.7 | 6.7, 12.8 | <0.001 | 12.8 | 10.6, 15.7 | <0.001 | 528 |
| | RES + RT | 9.0 | 1.4, 18.8 | 0.008 | 19.2 | 15.9, 21.5 | <0.001 | 107 |
| | RT + INT | 12.4 | -7.8, 28.0 | 0.178 | 13.3 | 1.8, 20.8 | 0.020 | 23 |
| | RES + INT | 11.7 | 1.4, 21.0 | 0.044 | 11.2 | 3.7, 20.1 | 0.005 | 28 |
| | RES + RT + INT | 20.1 | -1.3, 33.1 | 0.064 | 22.2 | 15.5, 36.1 | <0.001 | 12 |
| Soybean | RT | 3.0 | -16.3, 18.1 | 0.662 | 7.1 | -0.6, 12.9 | 0.067 | 47 |
| | INT | 13.0 | 1.1, 30.5 | 0.032 | 2.5 | -5.9, 11.7 | 0.625 | 39 |
| | RES | 12.3 | -25.7, 45.4 | 0.41 | 9.9 | 3.5, 15.7 | <0.001 | 36 |
| | RES + RT | 15.4 | -3.7, 47.4 | 0.153 | 13.9 | 7.4, 26.6 | 0.005 | 10 |
| | RT + INT | 20.2 | -18.1, 47.6 | 0.222 | 4.0 | -2.1, 15.4 | 0.211 | 12 |
| | RES + INT | 13.6 | 13.6, 13.6 | <0.001 | 0.8 | 0.8, 0.8 | <0.001 | 1 |
| | RES + RT + INT | NA | NA | NA | NA | NA | NA | 0 |
| Maize | RT | 4.1 | -3.9, 10.1 | 0.262 | 7.6 | -1.0, 12.9 | 0.102 | 190 |
| | INT | 13.1 | 0.9, 26.8 | 0.032 | -3.2 | -9.5, 4.0 | 0.348 | 152 |
| | RES | 15.2 | 10.1, 20.0 | <0.001 | 13.8 | 11.1, 17.9 | <0.001 | 537 |
| | RES + RT | 16.1 | 8.4, 21.7 | <0.001 | 18.6 | 13.3, 24.6 | <0.001 | 81 |
| | RT + INT | 1.3 | -16.1, 16.6 | 0.796 | 5.5 | -1.4, 14.2 | 0.123 | 34 |
| | RES + INT | 27.2 | 8.8, 62.0 | <0.001 | 14.7 | 7.8, 29.5 | <0.001 | 32 |
| | RES + RT + INT | 37.1 | 23.2, 54.8 | <0.001 | 28.2 | 23.0, 37.8 | <0.001 | 3 |

**Table S8 – Practice-specific *metaforest* variable importance.** Values refer to variable importance scores of variables that passed the 90% preselection threshold. Dashes indicate that those variables were either not assessed or did not pass the preselection threshold for that subset. Highlighted variables were considered important (Importance > 70) and were used to build partial regression models (Table S9).

| | | **Cover Cropping** | | **Crop Rotations** | | **Reduced Tillage** | | **Residue Retention** | |
|---|---|---|---|---|---|---|---|---|---|
| | | Yields | SOC | Yields | SOC | Yields | SOC | Yields | SOC |
| Climate | MAT | **100** | - | - | - | - | **74.4** | **100** | - |
| | Aridity | - | - | - | - | **84.1** | **87.1** | 64.9 | - |
| | PET | - | - | - | - | **100** | | **80.9** | - |
| | Elevation | - | - | - | - | - | - | **73** | - |
| | Climate Class | - | - | - | - | - | 51.5 | 58.5 | - |
| Soil | Drainage Class | 36.7 | - | - | - | - | - | - | - |
| | Topsoil Texture | 0 | - | 0 | - | 0 | - | - | - |
| | Background SOC | - | - | **95.4** | - | 54.4 | - | - | - |
| | pH | - | - | - | - | - | **100** | - | - |
| | CEC | - | - | - | - | 50.2 | - | - | **100** |
| | MAOC Saturation | - | - | - | - | - | 52.8 | - | - |
| | Median Slope | - | - | - | - | - | - | 9.8 | - |
| Farm Management | Crop N | **91.7** | **100** | **100** | - | - | - | **85** | - |
| | Crop Type | **70.5** | - | - | - | - | 0 | 5.2 | 34.8 |
| | Practice | - | - | - | **100** | - | 50.2 | 4.9 | 0 |
| | Irrigation | - | - | - | 0 | - | 5.0 | - | 17.1 |
| | Study Duration | - | - | - | - | - | 55.1 | 62.8 | **76.2** |
| | Yield Gap | - | - | - | - | - | - | 33.5 | - |
| | Residue Location | - | - | - | - | - | - | 0 | - |

**Table S9 – Models used for partial regressions.** Important variables (Importance > 70) in each of the practice subsets were combined to form a single model to predict yield and topsoil SOC outcomes across a gradient of one variable while holding other variables at their mean values. Partial regression models used *rma* function from the *metafor* package (Viechtbauer, 2010).

| **Practice** | **Model Formula** | **Predictive Performance ($R^2$)** | |
|---|---|---|---|
| | | Crop Yields | Topsoil SOC |
| Reduced Tillage | ln(RR) ~ PET + aridity + MAT + pH | 4.24% | 4.11% |
| Residue Retention | ln(RR) ~ PET + MAT + log(Elevation) + sqrt(CEC) + Crop N + log(Study Duration) | 4.8% | 6.64% |
| Cover Cropping | ln(RR) ~ MAT + Crop N + Crop Type | 28.92% | 11.47% |
| Crop Rotations | ln(RR) ~ sqrt(Background SOC) + Crop N + Practice | 9.76% | 17.72% |

**Table S10 – Variable importance values for different soil profiles.** As with Table S8, variable importance values here represent scores for variables that exceeded the preselection threshold. Highlighted variables were considered important (Importance > 70). Dashes indicate that those variables were either not assessed or did not pass the preselection threshold for that subset.

| | | **Subsoil SOC** | **Full Profile SOC** | **Microbial Biomass Carbon** |
|---|---|---|---|---|
| Climate | MAT | **90.2** | **86.2** | 38 |
| | Aridity | 59.6 | - | **100** |
| | Climate Class | - | - | 0.56 |
| Soil | Drainage Class | 42 | - | - |
| | Background SOC | 47.4 | - | - |
| | pH | - | **87.6** | - |
| | CEC | 23.1 | **100** | 24.7 |
| | MAOC Saturation | - | - | 49.7 |
| | Sand | - | - | 34 |
| | Bulk Density | - | - | 3.8 |
| | Topsoil SOC RR | - | - | 40.9 |
| Farm Management | Crop N | - | - | 0 |
| | Practice | **100** | - | 51.4 |
| | Irrigation | 0 | 0 | - |
| | Yield Gap | - | - | 0.75 |
| | Yield RR | - | 43.2 | - |

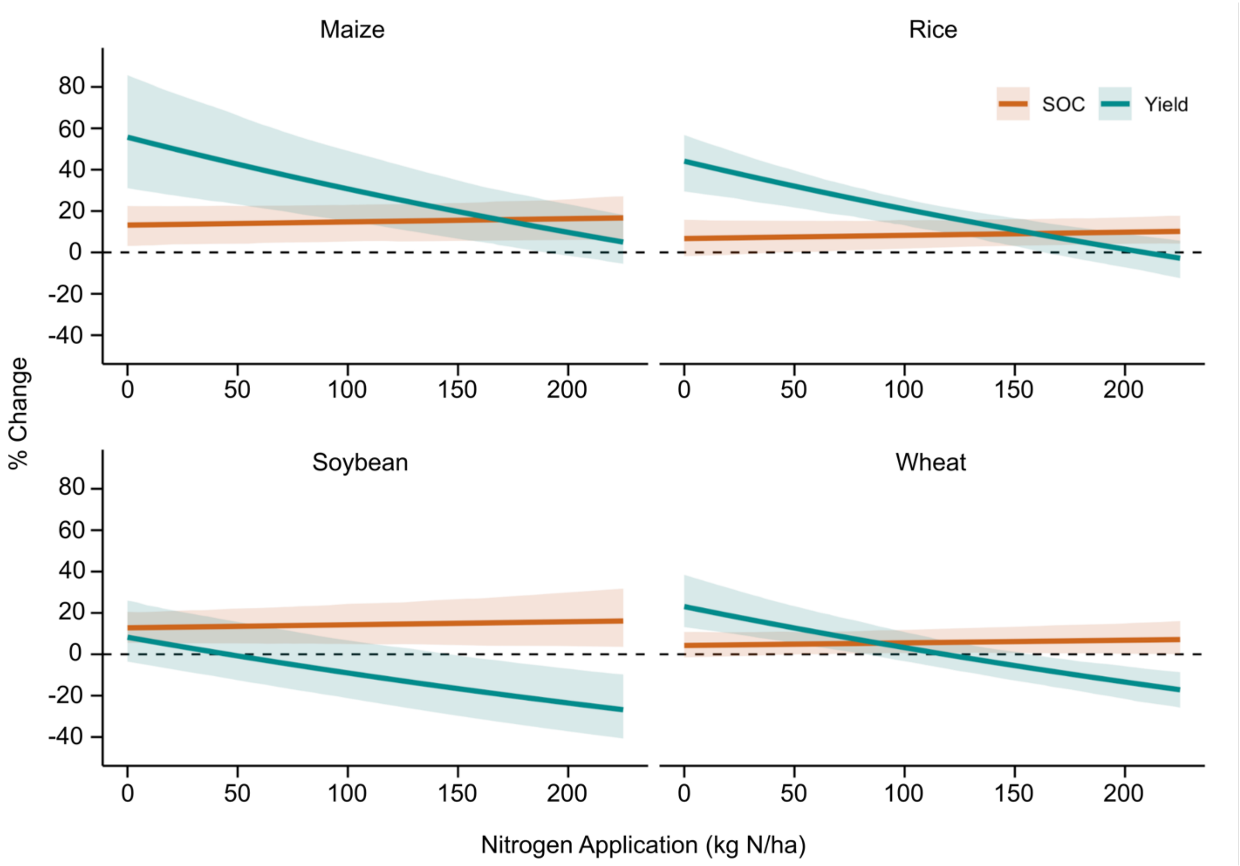


**Figure S5 – Cover crop nitrogen application rate partial regressions broken down by crop type**. See Figure 4 for description. Figure 4h shows effect of cover cropping across a gradient of nitrogen application rates, when the crop type is set to Rice. This figure shows how this relationship varies depending on the crop type. Prediction curves based on 100 equally spaced model predictions (see Supplementary Methods). Error bars represent bootstrapped 95% confidence intervals, such that if bars do not overlap the x-axis then the effects are significant.

**Table S11 – Effects of sustainable practices on SOC stocks in different soil profiles.** Shown graphically in Figure 6. Topsoil broadly defined as 0-30 cm, subsoil as >30 cm, and full profile encompassing observations that spanned the 30 cm threshold. Maximum depth in the full profile ranged from 35 cm to 105 cm. μ represents the mean value, 95% CI represents the 95% confidence interval of the 1000-iteration bootstrapped mixed effects models, and p represents the p-value. Topsoil, subsoil and full profile sample sizes are: Reduced tillage (RT: 532, 155, 158), Intensification (cover cropping and/or complex crop rotations) (INT: 494, 41, 44), Residue Retention (RES: 1458, 227, 243), Residue Retention and Reduced Tillage (RES + RT: 246, 43, 41), Reduced Tillage and Intensification (RT + INT: 69, 9, 9), Residue and Intensification (RES + INT: 131, 5, 17), all three practices (RES + RT + INT: 18, 6, 6).

| | **Topsoil** | | | **Subsoil** | | | **Full Profile** | | |
|---|---|---|---|---|---|---|---|---|---|
| Practice | μ | 95% CI | p | μ | 95% CI | p | μ | 95% CI | p |
| RT | 7.9 | 1.7, 11.9 | 0.006 | -24.3 | -40.0, 2.0 | 0.344 | -3.1 | -10.1, 8.2 | 0.690 |
| INT | 2.2 | -1.7, 6.5 | 0.264 | -7.3 | -32.1, 51.7 | 0.847 | -2.6 | -14.2, 20.1 | 0.789 |
| RES | 12.7 | 10.7, 15.5 | <0.001 | 43.7 | 4.0, 82.8 | <0.001 | 20.5 | 8.6, 29.0 | <0.001 |
| RES + RT | 18.4 | 13.1, 23.3 | <0.001 | -1.6 | -9.3, 5.3 | 0.626 | 17.2 | 11.1, 22.9 | <0.001 |
| RT + INT | 10.3 | 3.7, 17.2 | <0.001 | 10.5 | -23.9, 45.1 | 0.556 | 14.9 | -7.6, 29.9 | 0.218 |
| RES + INT | 11.9 | 6.8, 16.0 | <0.001 | 6.9 | -30.2, 48.3 | 0.419 | 12.9 | 4.3, 28.3 | <0.001 |
| RES + RT + INT | 21.4 | 16.1, 30.6 | <0.001 | 2.0 | -32.2, 36.3 | 0.606 | 17.4 | 3.9, 29.4 | 0.002 |

**Table S12 – Mixed effects model estimates**. Mixed effects models performed using the *rma.mv* function in the *metafor* package (Viechtbauer, 2010). Mean values and 95% confidence intervals are those used in Figure 2 and Figure 3. 95% CI represents the 95% confidence interval of the 1000-iteration bootstraped mixed effects models, p represents p-value, *n* represents the sample size. Note that Crop Type, Practice, Aridity Zone, Background SOC, MAOC Saturation, N Rate, and Study Duration are based on the full topsoil dataset. Cover crop functional group, residue location, and tillage type are based on practice-specific subsets (see section 1.5.1 for definition of practice-specific subsets). Only the "Intensification Type" category is divided into "Crop Rotation" and "Cover Crop" based on when either complex crop rotations or cover cropping are the sole practice difference between control and treatment groups. RT = Reduced Tillage, RES = Residue Retention, INT = Cropping Intensification (cover cropping and/or complex crop rotations). "+" denotes the stacking of multiple practices.

| | | Crop Yields | | | Topsoil SOC Stocks | | | |
|---|---|---|---|---|---|---|---|---|
| Variable | Level | Mean | 95% CI | p | Mean | 95% CI | p | *n* |
| Crop Type | Rice | 5.7 | -0.4, 12.0 | 0.074 | 10.0 | 8.6, 11.3 | <0.001 | 752 |
| | Wheat | 8.2 | 4.7, 12.3 | <0.001 | 10.6 | 9.5, 11.7 | <0.001 | 1038 |
| | Soybean | 12.9 | -3.7, 24.2 | 0.104 | 10.0 | 6.0, 13.6 | <0.001 | 145 |
| | Maize | 13.1 | 8.7, 17.7 | <0.001 | 10.5 | 9.4, 11.7 | <0.001 | 1040 |
| Practice | RT | 2.2 | -1.6, 5.4 | 0.216 | 7.9 | 1.7, 11.9 | 0.006 | 532 |
| | INT | 13.1 | 7.3, 19.7 | <0.001 | 2.2 | -1.7, 6.5 | 0.264 | 494 |
| | RES | 11.2 | 8.8, 13.7 | <0.001 | 12.7 | 10.7, 15.5 | <0.001 | 1458 |
| | RES + RT | 9.7 | 5.7, 13.2 | <0.001 | 18.4 | 13.1, 23.3 | <0.001 | 246 |
| | RT + INT | 8.9 | -1.4, 18.4 | 0.104 | 10.3 | 3.7, 17.2 | <0.001 | 69 |
| | RES + INT | 17.9 | 12.7, 23.5 | <0.001 | 11.9 | 6.8, 16.0 | <0.001 | 131 |
| | RES + RT + INT | 19.5 | 6.0, 25.8 | 0.014 | 21.4 | 16.1, 30.6 | <0.001 | 18 |
| Intensification Type | Crop Rotation | 0.4 | -10.7, 13.3 | 0.940 | 3.3 | 0.7, 6.6 | 0.024 | 233 |
| | Cover Crop | 26.6 | 13.1, 44.2 | <0.001 | 6.7 | 3.4, 9.8 | <0.001 | 246 |
| Cover Crop Functional Group | Brassica | 10.1 | -3.1, 22.9 | 0.132 | 8.4 | -0.7, 15.1 | 0.088 | 9 |
| | Grass | 10.1 | -6.3, 28.7 | 0.184 | 11.0 | 5.7, 17.1 | <0.001 | 45 |
| | Legume | 18.7 | 9.8, 27.7 | <0.001 | 9.0 | 5.9, 12.5 | <0.001 | 270 |
| | Mixture | 14.6 | 5.3, 24.4 | 0.012 | 11.6 | 6.6, 17.0 | <0.001 | 55 |
| Tillage Type | Minimum tillage | 5.3 | 2.3, 8.3 | <0.001 | 6.6 | 1.8, 10.6 | <0.001 | 235 |
| | No-tillage | 0.6 | -1.8, 3.0 | 0.624 | 11.4 | 9.1, 14.0 | <0.001 | 622 |
| Residue Location | Retained | 12.1 | 9.7, 14.5 | <0.001 | 10.8 | 7.4, 15.9 | <0.001 | 669 |
| | Incorporated | 12.5 | 10.4, 14.8 | <0.001 | 13.8 | 9.1, 17.4 | <0.001 | 939 |

| | | | | | | | | |
|---|---|---|---|---|---|---|---|---|
| Aridity Zone | Humid (AI >0.65) | 8.6 | 6.2, 10.8 | <0.001 | 9.6 | 8.4, 10.9 | <0.001 | 1200 |
| | Dry sub-humid (0.5 ≤ AI < 0.6) | 11.3 | 7.9, 14.9 | <0.001 | 12.6 | 9.4, 15.3 | <0.001 | 310 |
| | Semi-arid (0.2 ≤ AI < 0.5) | 10.6 | 8.3, 12.7 | <0.001 | 9.4 | 8.0, 10.9 | <0.001 | 1321 |
| | Arid 0 < AI < 0.2 | 4.2 | 0.0, 8.7 | 0.048 | 17.7 | 10.9, 24.1 | <0.001 | 182 |
| Background SOC | Low (<0.6%) | 10.5 | 7.0, 14.6 | <0.001 | 12.5 | 9.8, 17.0 | <0.001 | 466 |
| | Medium (0.6-1.2%) | 7.7 | 6.1, 9.3 | <0.001 | 10.2 | 9.0, 11.4 | <0.001 | 1815 |
| | High (>1.2%) | 12.9 | 8.8, 16.9 | <0.001 | 9.0 | 7.1, 10.9 | <0.001 | 686 |
| MAOC Saturation | Low (<20%) | 8.9 | 7.2, 10.5 | <0.001 | 11.0 | 9.9, 12.3 | <0.001 | 1803 |
| | Medium (20-40%) | 10.0 | 6.8, 13.1 | <0.001 | 8.6 | 6.9, 10.5 | <0.001 | 928 |
| | High (<40%) | 10.5 | 6.6, 17.2 | <0.001 | 10.6 | 6.2, 13.2 | <0.001 | 116 |
| N Rate | Unfertilized | 16.3 | 10.3, 23.3 | <0.001 | 10.9 | 5.7, 14.7 | <0.001 | 420 |
| | <160 kg N/ha | 11.8 | 7.5, 15.5 | <0.001 | 10.6 | 9.0, 12.5 | <0.001 | 1044 |
| | 160-240 kg N/ha | 6.0 | 1.8, 9.7 | 0.006 | 8.7 | 7.2, 10.3 | <0.001 | 634 |
| | >240 kg N/ha | 6.6 | 3.1, 11.1 | <0.001 | 11.1 | 9.2, 13.3 | <0.001 | 911 |
| Study Duration | <5 Years | 8.4 | 6.5, 10.7 | <0.001 | 7.4 | 5.3, 9.6 | <0.001 | 1692 |
| | 5-10 Years | 10.4 | 7.3, 12.9 | <0.001 | 12.3 | 9.1, 16.7 | <0.001 | 656 |
| | >10 Years | 10.9 | 6.8, 15.3 | <0.001 | 15.0 | 8.4, 19.7 | <0.001 | 665 |
